# Retinal bipolar cells borrow excitability from amacrine cells to amplify spatial features in dim light

**DOI:** 10.1101/2024.07.03.601922

**Authors:** Shubhash Chandra Yadav, Logan Ganzen, Lawrence Man, Hong Ma, Scott Nawy, Richard H Kramer

## Abstract

Bipolar cells (BCs) carry visual information from photoreceptors in the outer retina to retinal ganglion cells (RGCs) in the inner retina. Most BCs lack voltage-gated Na^+^ (NaV) channels and action potentials. Instead, voltage signals spread passively from input to output synapses. Despite this, we find that blocking NaV channels reduces synaptic output of On-cone bipolar cells (On-CBCs). These cells borrow NaV channel-mediated excitability from electrically coupled A2 amacrine cells to amplify excitatory output, allowing voltage changes to be transmitted more effectively to downstream RGCs in dim light. Because neighboring On-CBCs form electrical synapses onto the same A2 cell, a spatially adjacent group of On-CBCs is more effective than a dispersed set for recruiting A2 cell electrical excitability and driving post-synaptic RGCs. This mechanism filters the neural representation of an image to highlight shapes over spatially uncorrelated stimuli, a sophisticated feature detection property that is well-suited for object recognition.

## INTRODUCTION

Neurons in the brain use propagating action potentials to send signals over long distances. In contrast, many retinal neurons, including rod and cone photoreceptors, horizontal cells, and bipolar cells (BCs), are so short that action potentials are unnecessary. In BCs, graded voltage signals generated at input synapses in the outer retina spread passively to output synapses in the inner retina, eliciting graded neurotransmitter release. Consistent with the lack of action potentials, most mammalian BCs lack voltage-gated Na^+^ (NaV) channels**^1-7^**, and in the few BCs types where NaV channels are found, expression is insufficient to support spiking**^8,9^**.

Retinal ganglion cells (RGCs) and amacrine cells are the only classes of retinal neurons that do generate NaV-dependent action potentials. RGCs have axons that extend centimeters through the optic nerve to visual centers in the brain and NaV channels are needed for initiating and propagating spikes along these axons. Wide-field amacrine cells (wf-ACs) have long lateral processes projecting up to 2 mm, and in many wf-Acs, NaV channels support the propagation of dendritic spikes over these long distances**^10^**. Some medium-field amacrine cells (mf-ACs, typically 150-500 µm) are also excitable, including starburst amacrine cells that respond preferentially to visual stimuli moving in a particular direction**^11^**. But while dendritic excitability enhances directional selectivity of starburst cells**^12,13^**, it is not necessary for its occurrence**^14^**.

While many narrow field amacrine cells are inexcitable, some, such as the axon-less A2 amacrine cell, have abundant NaV channels, even though their dendritic processes extend only 30-40 µm. Rather than residing along dendrites, NaV channels in A2 cells cluster in a thin, unbranched appendage, devoid of synapses but rich in expression of ankyrin G, a NaV scaffolding protein normally found in the axon initial segment (AIS) of axon-containing neurons**^15^**. The role of NaV channels in A2 amacrine cells and the significance of this appendage, which is an electrotonic dead end, has remained a mystery.

In addition to long distance signaling, NaV-mediated excitability serves other functions. Because action potentials exhibit a threshold, NaV-mediated excitability can make dendritic integration of synaptic inputs a highly non-linear process, as has been observed in some wf-ACs**^16^**. Because action potentials are typically all-or none, NaV-mediated excitability can standardize depolarizing voltage changes, increasing the reliability of synaptic output. The rapid rate of rise of depolarization mediated by regenerative activation of NaV channels can accelerate synaptic output, as has been observed at the A2 output synapses**^17^**.

Our knowledge of AC function comes mostly from experiments utilizing natural light stimuli to activate rod and/or cone pathways. However, there are 15 types of BCs **^18,19^** and >60 types of ACs in the mouse retina, many of which converge onto common sets of RGCs**^20^**. This makes it difficult to evaluate the role of individual BC and AC types in determining RGC light responses and nearly impossible to understand how electrical excitability contributes. To simplify analysis, we targeted expression of an optogenetic tool to one specific type of On-CBC; the type 6 cone bipolar cell (CBC6), which forms an electrical synapse predominantly with one type of AC; the A2 amacrine cell**^21^**. In turn, the CBC6 cell makes a chemical excitatory synapse on the On-alpha RGC**^22^**, a ganglion cell type which plays a key role in processing both rod-mediated**^23^** and cone-driven signals**^24^**.

Studying this circuit in isolation is a challenge, but by pharmacologically blocking all inhibitory and excitatory chemical synaptic inputs onto CBC6 and A2 cells with neurotransmitter receptor inhibitors while sparing only the AMPA receptors on RGCs, we could selectively examine the effect of A2 cells on CBC6 cell output. Once we understood the role of NaV-mediated excitability in this isolated circuit, we could then evaluate its role in the intact retinal circuit responding naturally to cone-driven light stimuli. Using this strategy, we found that NaV-mediated excitability of A2 amacrine cells amplifies synaptic output of On-CBCs, but only to spatially clustered stimuli, providing a neural mechanism for distinguishing objects over spatially distributed background, early in the visual system.

## RESULTS

### Investigating voltage-gated Na^+^ channels and intrinsic excitability of On-CBCs and A2 amacrine cells

To investigate the possible presence of voltage-gated Na^+^ channels in On-CBCs and A2 amacrine cells, we recorded voltage-gated ionic currents from these cell types under whole-cell patch clamp. To isolate voltage-gated currents explicitly from the recorded cells, the retinas were pre-treated with 100 µM meclofenamic acid (MFA), an uncoupler of gap junctions**^25^**. First, we investigated CBC6 cells in the whole-mount mouse retinas. To identify and record from CBC6 cells, we utilized a cre-driver mouse line, enabling transgene expression of a fluorescent protein marker, eYFP, in CBC6 cells, but no other types of BCs**^26^**. eYFP was fused to channelrhodopsin-2 (ChR2), which we used for optogenetic manipulation in later experiments. The pipette solution contained agents that blocked voltage-gated K^+^ channels and the extracellular solution had agents that blocked current through V-gated Ca^2+^ channels, allowing measurement of NaV current in isolation.

In CBC6 cells, depolarizing voltage steps from a holding potential of -70 mV to 0 mV elicited almost no inward current and bath application of the NaV channel blocker tetrodotoxin (TTX) had no effect (Fig. 1a), consistent with a previous study suggesting that NaV channels are absent from CBC6 cells**^6^**. We next obtained retinal slices and recorded from A2 amacrine cells, which were tentatively identified by their size and laminar location. The identity of putative A2 cells was verified by fluorescent dye-filling to reveal their distinctive axodendritic morphology. In verified A2 cells, the depolarizing voltage steps above -40 mV evoked a large inward current, which was blocked completely by TTX (Fig. 1b) consistent with previous studies showing expression of NaV channels and firing of action potentials**^27,28^**. Current-voltage curves substantiated that while CBC6 cells lack NaV current, A2 amacrine cells have NaV current that can be blocked with TTX (Extended Data Fig. 1a, b).

**Figure 1.**
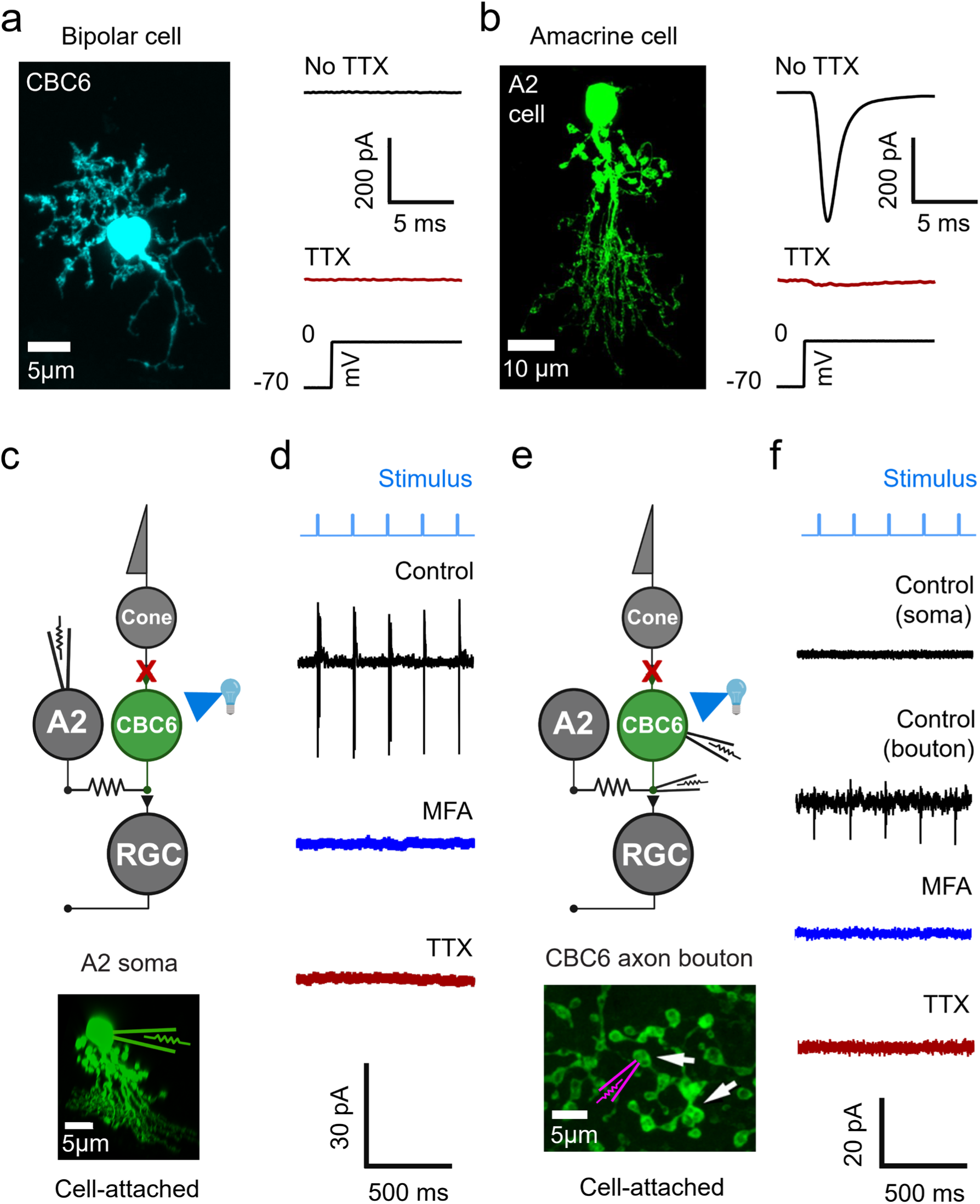
A2 amacrine cells exhibit NaV channel-mediated excitability but CBC6 bipolar cells do not. **a,b)** Intrinsic electrical properties of an A2 cell and an CBC6 cell, evaluated with the whole-cell patch clamp technique. Cells were filled with fluorescent dye (Alexa-594) and imaged with confocal microscopy to aid in visual identification. Under voltage-clamp, the A2 cell (b) exhibits depolarization-gated transient inward current that was blocked by TTX, characteristic of NaV channels, whereas the CBC6 cell (a) showed no inward current and no effect of TTX. **c-f)** Extracellular cell-attached patch recordings from A2 or CBC6 cells in a retina where CBC6 cells express the optogenetic tool ChR2, as depicted in the retinal circuit diagrams (green). Brief flashes of light (10 ms) elicited spikes in the soma of the A2 cell **(c-d)**, but not in the soma of the CBC6 cell **(e-f)**. However, recording from the CBC6 axon terminal did reveal small spikes. Spikes in both A2 somata and CBC6 terminals were eliminated with TTX, implicating NaV channels. Spikes were also eliminated by treatment with MFA, implicating gap junctions in transmitting the optogenetic depolarization from the CBC6 terminal to the A2 cell and evoked spikes from the A2 cell back to the CBC6 terminals. See Extended Data Fig. 1.

We next obtained cell-attached recordings on CBC6 cells or A2 cells in whole-mount retinas to determine whether optogenetic stimulation can elicit action potentials in either of these cells (Fig. 1c, e). To eliminate light responses transmitted from rod and cone photoreceptors, synaptic transmission was blocked with agonist of mGluR6 receptors on On-BCs, and antagonist of kainate receptors on Off-BCs leaving ChR2 in the CBC6 cell as the only relevant light sensor. A2 somata were targeted with the help of mCherry expression (see Methods). Recordings from A2 somata showed consistent all-or none transient spikes to each light flashes (Fig. 1d). TTX eliminated these events, consistent with NaV-mediated action potentials. Recordings from CBC6 cell somata showed no detectable spikes; however, recordings from presynaptic boutons of CBC6 cells revealed small spikes that were eliminated by adding TTX (Fig. 1f).

In addition to having an output synapse onto RGCs, CBC6 cells connect with A2 amacrine cells via gap junctions, localized on the CBC6 synaptic terminal in the On-lamina of the inner plexiform layer (IPL)**^29^**. We hypothesized that the spikes detected in CBC6 cell terminals arose in the A2 cell and spread into the CBC6 cell through these gap junctions. To test this, before recording we pre-treated whole-mount retinas for 45 minutes with MFA. Under these conditions, action potentials were absent from both CBC6 cell terminals and A2 cells (Extended Data Fig. 1c, d). Together, these results indicate that even though CBC6 cells do not possess their own NaV channels, their voltage response is influenced by NaV channels in their electrically coupled partner, the A2 amacrine cell.

### NaV channels amplify synaptic transmission from CBC6 cells

To evaluate chemical synaptic output from CBC6 cells, we stimulated the cells optogenetically and measured responses in a postsynaptic On-alpha RGC (Fig. 2a). Once again, synaptic inputs from rods and cones were blocked pharmacologically. Brief flashes of light (2 ms) activated ChR2 in the CBC6 cells, depolarizing the membrane potential enough to activate voltage-gated Ca^2+^ channels in their terminals, leading to release of neurotransmitter onto RGCs**^30^**. Excitatory postsynaptic currents (EPSCs) in RGCs reflect simultaneous stimulation of hundreds of CBC6 cells converging onto an RGC, explaining its large amplitude (up to 3 nA). We found that TTX reduced the peak amplitude of the EPSC by almost 70% (control=1894 ± 185 pA; TTX=574 ± 99 pA; n=16 each; p<0.0001) (Fig. 2b, d). This indicates that synaptic release from CBC6 is amplified by NaV channels, presumably in A2 amacrine cells since the CBC6 cell have none of their own. TTX reduced the EPSCs across a wide range of optogenetic stimulus strengths (Fig. 2h), indicating that NaV channels serve to increase the gain of CBC6 cell to RGC synaptic transmission. TTX also increased the rise time of the EPSC (control=9.8 ± 0.4 ms; TTX=13.4 ± 0.6 ms; n=16 each; p<0.0001) and slowed its decay time (control=39.2 ± 4.7 ms; TTX=83.4 ± 12.1 ms; n=16 each; p=0.0029) (Fig. 2b, e).

**Figure 2.**
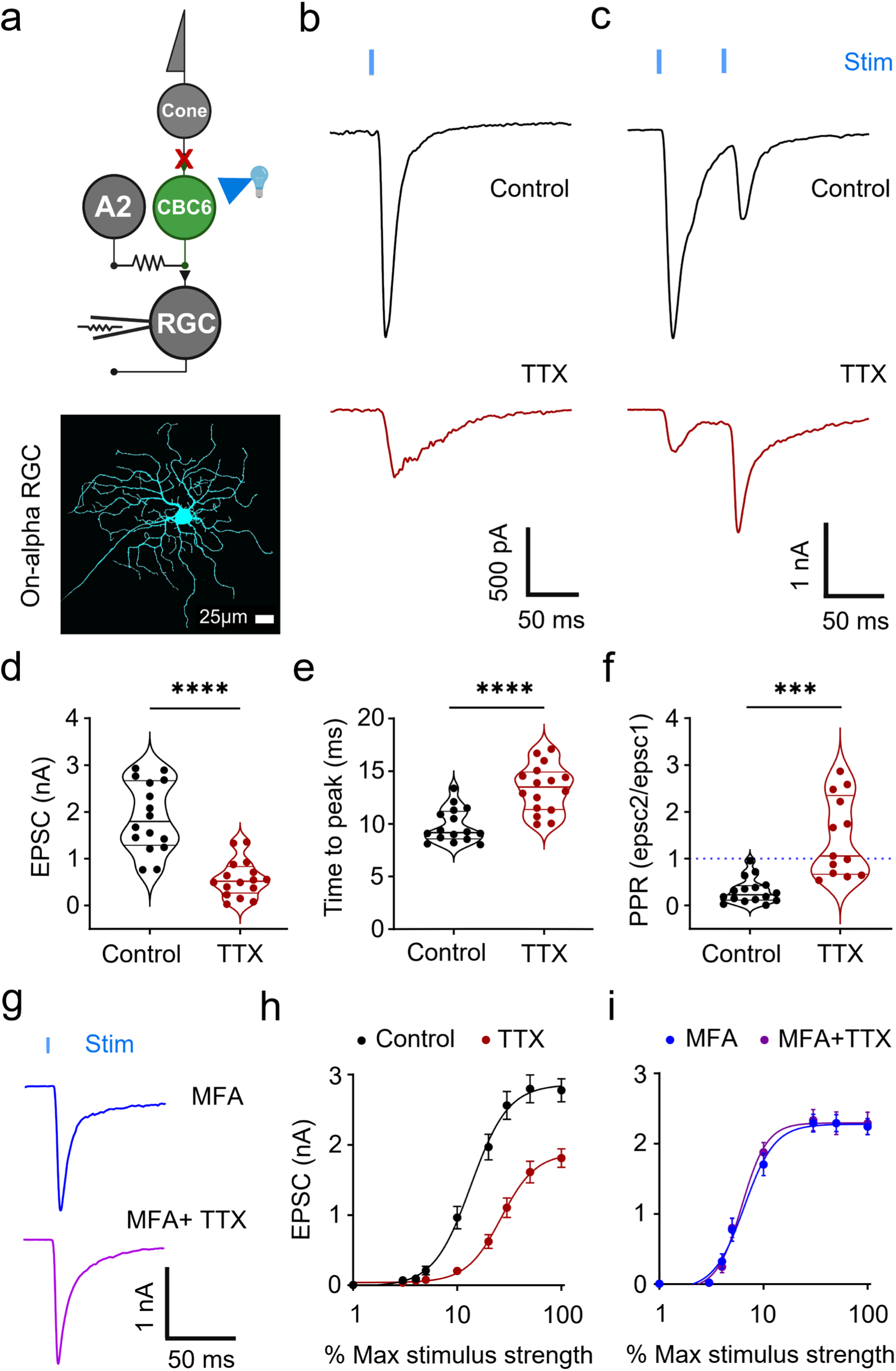
NaV channel-mediated excitability, spreading through gap junctions, amplifies synaptic output of CBC6 cells. **a)** Retinal circuit diagram showing the experimental arrangement. Full-field light flashes were used to optogenetically stimulate CBC6 cells and whole-cell patch clamp recordings were obtained from On-alpha RGCs to measure evoked EPSCs. **b)** EPSCs elicited by a single flash (2 ms) were reduced in amplitude and slowed by adding TTX. **c)** EPSCs elicited by a pair of flashes (inter-stimulus interval=50ms) decremented without TTX but grew in amplitude with TTX. **d-e)** Violin plots showing group data for **d)** amplitude and **e)** time-to-peak of EPSCs elicited by a single stimulus without or with TTX. Note that TTX reduced the EPSC amplitude by almost 70% (control=1894 ± 185 pA; TTX=574 ± 99 pA; n=16 each; p<0.0001); unpaired t-test), and slowed the peak by almost 50% (control=9.8 ± 0.4 ms; TTX=13.4 ± 0.6 ms; n=16 each; p<0.0001; unpaired t-test). (**f)** Group data for paired-pulse ratio (PPR) of EPSCs elicited by pairs of stimuli without or with TTX. Note that TTX changed PPR from depressing to facilitating (PPR: control=0.31 ± 0.07, n=16; TTX=1.46 ± 0.23, n=13; p=0.0003; unpaired t-test). **g)** Pre-treatment with MFA to uncouple gap junctions eliminated the effect of TTX on EPSCs, consistent with removal of the A2 amplifier. **h)** The stimulus-response relationship of the CBC6 to RGC synapse, obtained by varying optogenetic stimulus strength and measuring evoked EPSCs. In TTX, stronger stimuli were required to evoke equivalent responses and saturating EPSC amplitude was decreased (two-way ANOVA; treatment × stimulus strength: F(2,69)=19.31; p˂0.0001; control, n=16; TTX, n=18). **i)** Pre-treatment with MFA eliminated the effect of TTX on CBC6 synaptic output (two-way ANOVA; treatment × stimulus strength: F(2,41)=0.2795; p=0.7866; No TTX, n=11; TTX, n=12). Violin plots depict median, 25^th^ and 75^th^ quartiles. Data in h, i are mean ± sem.

Strong or repeated depolarization of presynaptic neurons can lead to depletion of the readily-releasable pool (RRP) of synaptic vesicles, resulting in synaptic depression**^31^**. We found that pairs of optogenetic stimuli, separated by an interval of 50 ms, leads to paired-pulse depression of EPSCs recorded in RGCs, with the second being much smaller than the first (Extended Data Fig. 2a). However, addition of TTX reduced paired-pulse depression, usually replacing it with paired-pulse facilitation (Fig. 2c, f), where the second stimulus elicits a larger response than the first (Extended Data Fig. 2b).

A possible explanation for all of these findings is that NaV channel-mediated excitability of A2 cells amplifies and accelerates the depolarization of the CBC6 terminal, leading to increased activation of voltage-gated Ca^2+^ channels, increased Ca^2+^ entry, and enhanced Ca^2+^-dependent fusion of synaptic vesicles, resulting in larger and faster EPSCs**^32^**. If the amplification were sufficiently large to deplete the RRP, release elicited by a second stimulus would be reduced, contributing to paired-pulse depression. In addition, if the excitability of A2 cells exhibited refractoriness, perhaps because of NaV inactivation, a second pulse might fail to generate an amplified response. Central to all of these hypotheses is the role of gap junctions in enabling the initial depolarization to spread from CBC6 cells to A2 cells and allowing the amplified response to spread back into the CBC6 terminals.

To test the role of gap junctions in enabling amplification, we treated whole-mount retinas with MFA. Under these conditions, when gap junctions were uncoupled, ESPCs were unaffected by TTX (Fig. 2g). In fact, while TTX normally reduced CBC6 synaptic transmission (Fig. 2h), TTX had no effect on EPSCs over a wide range of stimulus strengths after MFA treatment (Fig. 2i). These findings are consistent with NaV-mediated amplification arising from A2 cells, which MFA had removed from influencing CBC6 cell synaptic release.

In addition to A2 amacrine cells, which have short dendritic projections and respond to narrow-field illumination, the retina has many types of wf-ACs that also express NaV channels**^33^**. Some wf-ACs receive direct or polysynaptic inputs from CBC6 cells and they, in turn, synapse onto On-alpha RGCs**^34,35^**.

This raises the possibility that the effects of TTX on CBC6 synaptic output might involve NaV channels in one or more wf-ACs, rather than, or in addition to, A2 cells. To test this possibility, we blocked chemical synaptic inhibition with antagonists of glycine, GABA_A_ and GABA_C_ receptors. We found that neither the peak optogenetic EPSC nor its rise time were altered under these conditions (Extended Data Fig. 3b, c). Blocking inhibition did slow the decay of the EPSC (Extended Data Fig. 3b, d), consistent with the polysynaptic wf-AC pathway exerting feedback inhibition onto bipolar cell terminals with a delay**^36^**. TTX reduced the peak amplitude of the EPSC to the same extent whether or not chemical inhibition was blocked (Extended Data Fig. 4). These results suggest that amplification of rapid synaptic transmission from CBC6 cells to On-alpha RGCs involves A2 amacrine cells rather than wf-ACs.

To further verify the role of NaV channels in A2 cells as amplifiers of CBC6 synaptic output we used QX-314, an open-channel blocker of the pore of voltage-gated ion channels, including NaVs. As before, we optogenetically stimulated CBC6 cells and recorded EPSCs in an On-alpha RGC, but we conducted this experiment in retinal slices (200 µm), to reduce the number of CBC6 cells and A2 cells involved in generating the RGC response.

First, we obtained a baseline EPSC (Fig. 3a, left). Then we impaled an A2 cell with a sharp pipette containing QX-314 (1 mM) together with the fluorescent dye Alexa 594.

**Figure 3.**
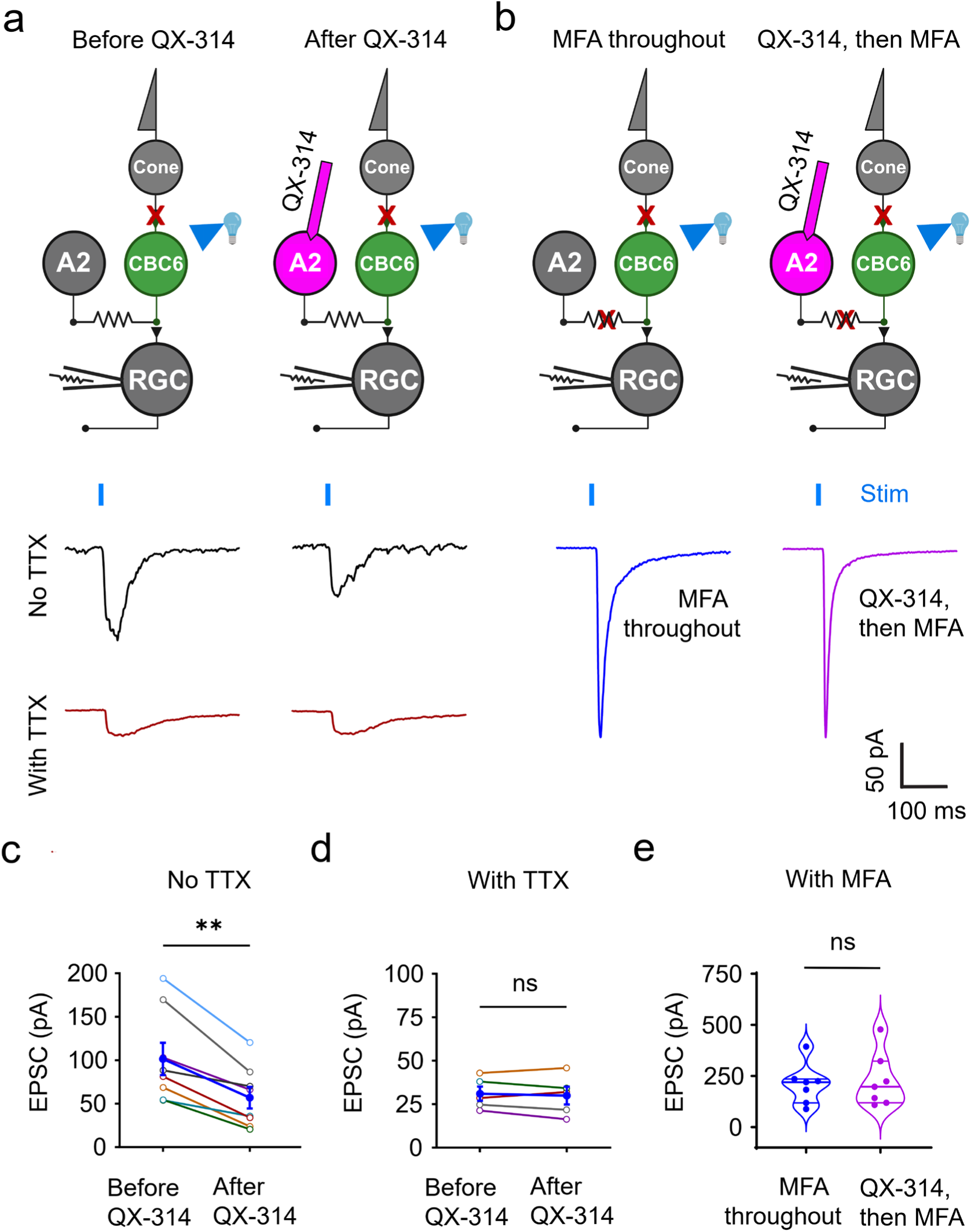
Contribution of a single A2 cell to amplifying CBC6 synaptic output. **a)** Top: Retinal circuit diagram showing experimental arrangement. CBC6 cells were optogenetically stimulated in a retinal slice, eliciting EPSCs in RGCs, before and after injecting QX-314 into a single A2 amacrine cell with a sharp electrode. Bottom: Without TTX in the bathing solution, EPSCs in the RGC were reduced following QX-314 injection. However, with TTX in the bathing solution, QX-314 injection had no effect. **b)** Top: Retinal circuit diagram, showing experimental arrangement. To control for possible diffusion of QX-314 from the A2 cell into the CBC6 cell, at 10 min after QX-314 injection, MFA was applied to uncouple gap junctions connecting the A2 cell to CBC6 cells. Bottom: EPSCs elicited after QX-314 injection followed by MFA treatment did not differ from EPSCs elicited in MFA-treated slices without QX-314 injection, suggesting no significant effect of QX-314 that might have entered CBC6 cells. **c)** Without TTX, QX-314 injection reduced EPSC amplitude (before QX-314 = 102 ± 19 pA; after QX-314=57 ± 12 pA; n=8 each, p=0.0011; paired t-test). **d)** With TTX, QX-314 injection had no effect (before QX-314=31 ± 4 pA; after QX-314=30 ± 5 pA; n=5 each, p=0.5723, paired t-test). Each thin line represents the same RGC recording, before and after QX-314, The thick black line represents mean responses across all RGCs. **e)** Group data indicate that QX-314 injection into A2 cells is inconsequential when gap junctions are uncoupled (before QX-314=227 ± 50 pA; after QX-314=207 ± 37 pA, n=7 each, p=0.7527, unpaired t-test).

In every case, the amplitude of the EPSC was decreased significantly after introducing QX-314 as compared to before (Fig. 3c). QX-314 injection had no effect on the EPSC in control experiments in which TTX was present throughout the experiment (Fig. 3a, d). We were concerned that some of the QX-314 injected into the A2 cell had diffused through the gap junctions into the CBC6 terminal, where it might advertently block their voltage-gated channels, including Ca^2+^ channels, thereby reducing evoked neurotransmitter release apart from affecting NaV channels.

To exclude this possibility, we measured optogenetic EPSCs before and after QX-314 injection, and then again, after treating with MFA to uncouple the gap junctions (Fig. 3b). If QX-314 had crossed from the A2 cell into the CBC6 terminal, the evoked EPSC would have been smaller than if QX-314 had not been injected into the A2 cell in the first place. However, we observed no difference in EPSC amplitude elicited by activating uncoupled CBC6 cells, whether an A2 had received QX-314 or not (Fig. 3e), inconsistent with local channel blockade within a CBC6 cell. Taken together, these experiments definitely identify the A2 cell as the source of NaV-mediated excitability that amplifies CBC6 synaptic transmission to RGCs.

By removing NaV-mediated amplification, one might expect that uncoupling CBC6 cells from A2 cells would reduce the amplitude of EPSCs recorded in the RGCs. However, we found that EPSC amplitude either increased or remained unaltered after treatment with MFA (Fig. 2, 3). Consistent with a previous study**^37^**, our experiments revealed that gap junctions set the resting membrane resistance of On-CBCs including CBC6 cells, and that depolarization, whether elicited by physiological or optogenetic stimuli, is reduced by this gap junctional shunt (Supplementary information). Activation of NaV channels in the A2 cell helps offset this signal reduction, ensuring that depolarization of CBC6 terminals can effectively activate voltage-gated Ca^2+^ channels and trigger neurotransmitter release.

### Consequences of the A2 amplifier for spatial processing

The mouse retina exhibits a high degree of convergence, with ∼6.7 million rods and cones generating signals carried to the brain by ∼50,000 RGCs**^38^**. Within the cone CBC6 subcircuit, 2-3 cones converge onto each CBC6 cell dendritic tree**^39^**, ∼10 CBC6 cells normally converge onto each A2 amacrine cell**^21,40^**, 5-10 A2s likely converge onto a CBC6 cell, and 200-400 CBC6 cells converge on each On-alpha RGC**^22^**. We evaluated how this circuit arrangement, combined with NaV-mediated A2 cell excitability, influences spatial encoding by the circuit.

We started by optogenetically activating an array of CBC6 bipolar cells with a bar of light while measuring the resulting compound EPSPs in an A2 amacrine cell (Fig. 4a). The A2 cell was identified with the help of mCherry expression and visualized by dye-filling with Alexa 594, which also enabled accurate positioning of the stimulus. We used the same mixture of neurotransmitter receptor inhibitors described above to prevent natural light responses of rods and cones from influencing the downstream circuity.

**Figure 4.**
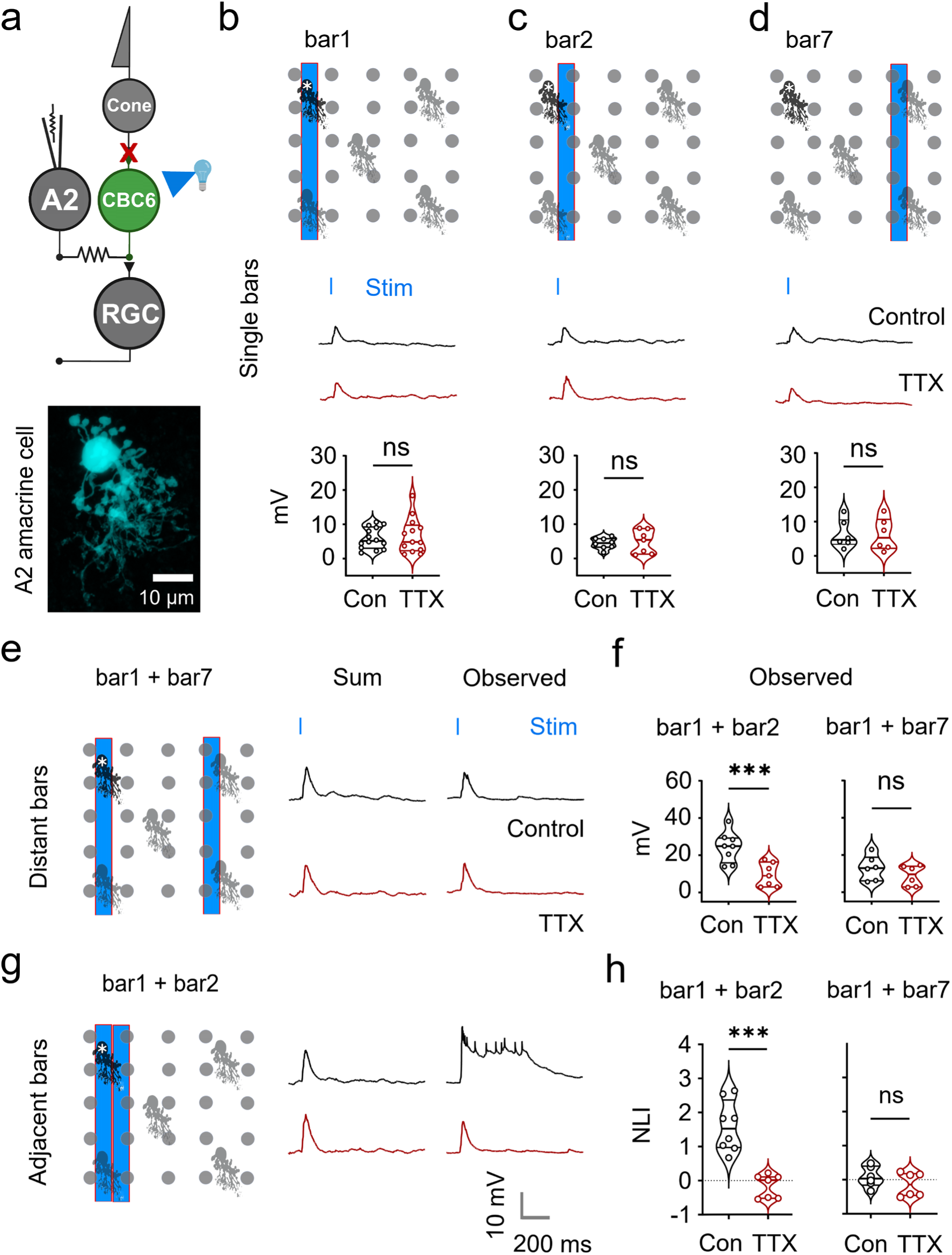
NaV-mediated excitability in A2 cells results in supra-linear summation of voltage responses driven by spatially adjacent CBC6 cells. **a)** Retinal circuit diagram, showing experimental arrangement. **b-d)** A narrow bar of light (18 µm wide, 360 µm long) was projected onto the retina to optogenetically stimulate rows of CBC6 cells at three different positions relative to the patch-clamped A2 cell; **b)** bar 1: immediately over the A2 cell (asterisk), **c)** bar 2: 18 µm off-center from the A2 cell, or **d)** bar 7: 90 µm off-center from the A2 cell. Membrane potential of the A2 cell was allowed to “float” (i.e., not voltage-clamped) to enable recording of EPSPs and associated regenerative depolarizations. Bottom panel shows group data and variability of EPSP amplitudes evoked by each bar. Note that the amplitudes were nearly the same at all three positions, and that TTX had no significant effect (bar1: control=5.89 ± 0.83 mV, n=14; TTX=6.66 ± 1.37 mV, n=13; p=0.6441; bar2: control=4.42 ± 0.42 mV, n=8; TTX=4.86 ± 0.90 mV, n=7; p=0.7597; bar7: control=6.29 ± 1.11 mV, n=6; TTX=6.17 ± 1.26 mV, n=6; p=0.9653; unpaired t-test in each case). **e, g).** Results elicited by projecting pairs of bars on the flat-mount retinas. Two bars of light, each 18 µm wide as in **b-d**, were separated either by 90 µm (distant) or 0 µm (adjacent). The observed EPSPs for each is shown in comparison to the predicted EPSP, calculated assuming linear summation of two of the single bar EPSP responses shown in panels b-d. For distant bars, the observed peak response was similar to the summed response and neither were affected by TTX. However, for adjacent bars, the observed peak response was larger than the predicted summed response and it was followed by a prolonged plateau potential with small spikes. **f)** Group data for each A2 cell recording, as depicted by the representative responses in e and g. Distant bars (bar1 + bar7) produced EPSPs almost twice as a single bar, which were unaffected by TTX (control=13.02 ± 1.77 mV, TTX=8.84 ± 1.47 mV, n=6 each; p=0.2629; unpaired t-test). Adjacent bars (bar1 + bar2) produced 2-3-fold larger EPSPs than predicted (observed=24.33 ± 2.17 mV, predicted=9.91 ± 1.00 mV, n=8 each; p=0.0004; paired t-test), which was reduced substantially in the presence of TTX (observed: control= 24.33 ± 2.17 mV, n=8; TTX=9.45 ± 1.68 mV, n=7; p=0.0016; unpaired t-test). **h)** Group data of the non-linearity index (NLI) of A2 cell responses in response to simultaneous presentation of distant or adjacent bars. Distant bars produced either linear or sublinear responses irrespective of TTX application (control=0.08 ± 0.08, TTX= -0.14 ± 0.09; p= 0.3939; Mann-Whitney test). Adjacent bars produced supralinear responses, which were abolished by TTX (control=1.59 ± 0.20, TTX= -0.16 ± 0.09; p=0.0003; Mann-Whitney test). Non linearity index was calculated as follows: EPSP (Observed – Predicted)/Predicted. Values above and below 0 indicate supralinear and sublinear responses, respectively.

A narrow bar of light (18 µm) positioned either over the axodendritic tree of the A2 cell or 90µm away from the A2 somata generated small EPSPs, which were unaffected by adding TTX (control=5.55 ± 0.80 mV, n=14; TTX=6.07 ± 1.25 mV, n=13; p=0.9076). TTX did not alter the response of A2 cells, irrespective of the location of the bar of light (Fig. 4b-d). Stimulation with two bars of light positioned far apart (90 µm) generated an EPSP that was roughly twice as large as a single bar and again, the response was unaffected by TTX (control=13.02 ± 1.77 mV; TTX=8.84 ± 1.47 mV; n=6 each; p=0.2629) (Fig. 4e, f). However, positioning the two bars immediately adjacent to one another resulted in an EPSP that was 4-5-fold larger than from a single bar and almost 2-3-fold larger than predicted from linearly summing the responses (predicted=9.91 ± 1.00 mV; observed=24.33 ± 2.17 mV; n=8 each; p=0.0004).

Moreover, the EPSP was much longer-lasting (>than 500 msec), and it was followed by a plateau potential with a train of small spikelets on top (Fig. 4g, h). Addition of TTX eliminated the spikelets and the plateau potential, and brought the EPSP amplitude in agreement with that predicted from linear summation (predicted=11.39 ± 1.49 mV; observed=9.45 ± 1.68 mV; n=7 each; p=0.2497). Taken together, these results suggest that NaV-mediated excitability of A2 cells leads to non-linear summation of synaptic inputs from multiple CBC6 cells, if those CBC6 cells are close enough to converge onto the same A2 cell.

Next, we asked whether the preference for spatially adjacent stimuli, activating neighboring CBC6 cells, also applies to RGCs. We used the same bars of light to stimulate CBC6 cells, but recorded EPSCs in On-alpha RGCs under voltage-clamp (Fig. 5a). Individual bars of light positioned at various locations with respect to the dendritic tree of an RGC generated similar EPSCs and TTX had no significant effect (control=151 ± 10 pA, n=7; TTX=171 ± 9.7 pA, n=8; p=0.0891). TTX did not alter the response of On-alpha RGCs irrespective of the location of single bars (Fig. 5b-d). Two bars, positioned far apart (90µm) resulted in EPSCs that were roughly twice as large, which were unaffected by adding TTX (control=239 ± 27 pA, n=7; TTX=286 ± 32 pA, n=8; p=0.2789) (Fig. 5e). However, two adjacent bars of light resulted in EPSCs that were nearly 2-fold larger than predicted (predicted=322 ± 22 pA; observed=614 ± 48 pA; n=7 each; p=0.0156). Once again, the non-linear summation of the response was eliminated by adding TTX (predicted=351 ± 26 pA; observed=333 ± 29 pA; n=8 each; p=0.4916) (Fig. 5f-g), implicating NaV channels in A2 amacrine cell as the critical source.

**Figure 5.**
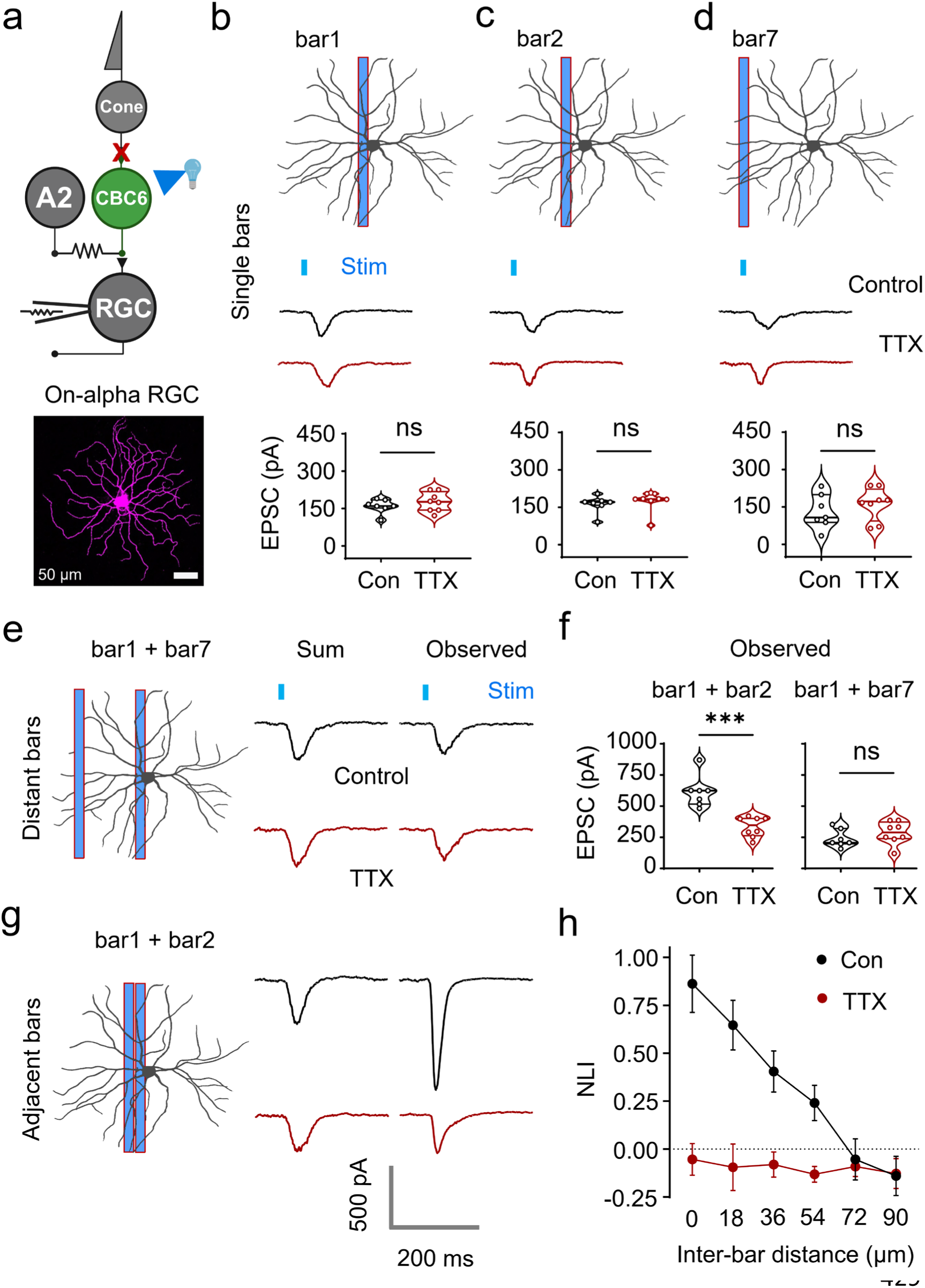
NaV-mediated excitability amplifies optogenetically-elicited synaptic output from spatially adjacent CBC6 cells to an individual RGC. **a)** Retinal circuit diagram, showing experimental arrangement. b-d) A narrow bar of light (18 µm wide, 360 µm long) was projected onto the retina to optogenetically stimulate rows of CBC6 cells at three different positions relative to the patch-clamped On-alpha RGC; b) bar 1: immediately adjacent to the RGC soma, c) bar 2 : 18 µm off-center from the RGC soma, or d) bar 7: 90 µm off-center from the RGC soma. Bottom panel shows group data and variability of EPSC amplitudes evoked by each bar. Note that the amplitude was nearly the same at all three positions, and that TTX had no significant effect (bar1: control=160 ± 10 pA, n=7; TTX=176 ± 14 pA, n=8; p=0.3562; unpaired t-test; bar2: control=163 ± 12 pA, n=7; TTX=174 ± 14 pA, n=8; p=0.0693; Mann-Whitney test; bar7: control=131 ± 24 pA, n=7; TTX=162 ± 23 pA, n=8; p=0.3932; unpaired t-test) **e, g).** Results elicited by projecting pairs of bars onto the retina. Two bars of light, each 18 µm wide as in **b-d**, were separated either by 90 µm (distant) or 0 µm (adjacent). The observed EPSCs for each is shown in comparison to the predicted EPSC, calculated assuming linear summation of two of the single bar EPSC responses shown in panels b-d. **f)** Group data for each On-alpha RGC recording, as depicted by the representative responses in e and g. Distant bars (bar1 + bar7) produced EPSCs almost twice as a single bar, which were unaffected by TTX (control=239 ± 27 pA, n =7; TTX=286 ± 32 pA, n=8; p=0.2789; unpaired t-test). Adjacent bars (bar1 + bar2) produced almost 2-fold larger EPSCs than predicted (observed=614 ± 48 pA, predicted=322 ± 22 pA, n=7 each; p=0.0156; Wilcoxon signed rank test), which was reduced substantially in the presence of TTX (observed: control=614 ± 48 pA, n=7; TTX=333 ± 29 pA, n=8; p=0.0005; unpaired t-test). **h)** Group data showing the non-linearity index (NLI) of On-alpha RGC responses elicited by bars separated by different distances. Note that the NLI was minimal at separations ≥ 72 µm, maximal for adjacent bars and half-maximal at ∼36 µm. The observed supralinear responses were abolished in the presence of TTX (two-way ANOVA; treatment × bar separation: F(2, 25)=11.51; p=0.0002; control, n=6; TTX, n=7). Data in h are mean ± sem.

A2 amacrine cells have a dendritic tree that is roughly 36 µm in diameter. However, A2 cells not only couple to CBC6 cells through gap junctions, they are also coupled by gap junctions to one another. To determine the minimum bar spacing necessary to generate supra-linear summation measured in RGCs, the distance between the bars was narrowed progressively until they were adjacent. We found that the response to bars of light increased supra-linearly at inter-bar distances <72 µm, with a half-maximal effect at about 36 µm (Fig. 5h). This suggest that simultaneously activated CBC6 inputs are most likely to be amplified if the inputs converge on a single A2 cell of 36 µm diameter, although inputs onto a neighboring A2 cell also contribute to the amplification, but to a lesser extent. Taken together, these findings suggest that spatial summation of excitatory inputs from CBC6 cells onto a single A2 cell is necessary to bring the cell above action potential threshold, thereby triggering amplification of synaptic transmission from CBC6 cells to RGCs.

### A2 amacrine-cell mediated amplification applies to natural light responses triggered by cone photoreceptors

Up until this point, we have relied exclusively on optogenetic stimulation of CBC6 cells to elicit light responses in A2 amacrine cells and On-alpha RGCs. We wondered whether A2 cell amplification also applies to light responses triggered naturally by cone photoreceptors (Fig. 6a). To address this question, we omitted all of the pharmacological blockers used in our other experiments, and bathed the retina in normal physiological saline. Dissection of the retinas was performed under photopic light conditions that presumably photobleached or saturated rods, eliminating their light responses**^41,42^**. Thereafter, these experiments were carried out in mesopic background light following the recovery of cone photoreceptors (see Methods).

**Figure 6.**
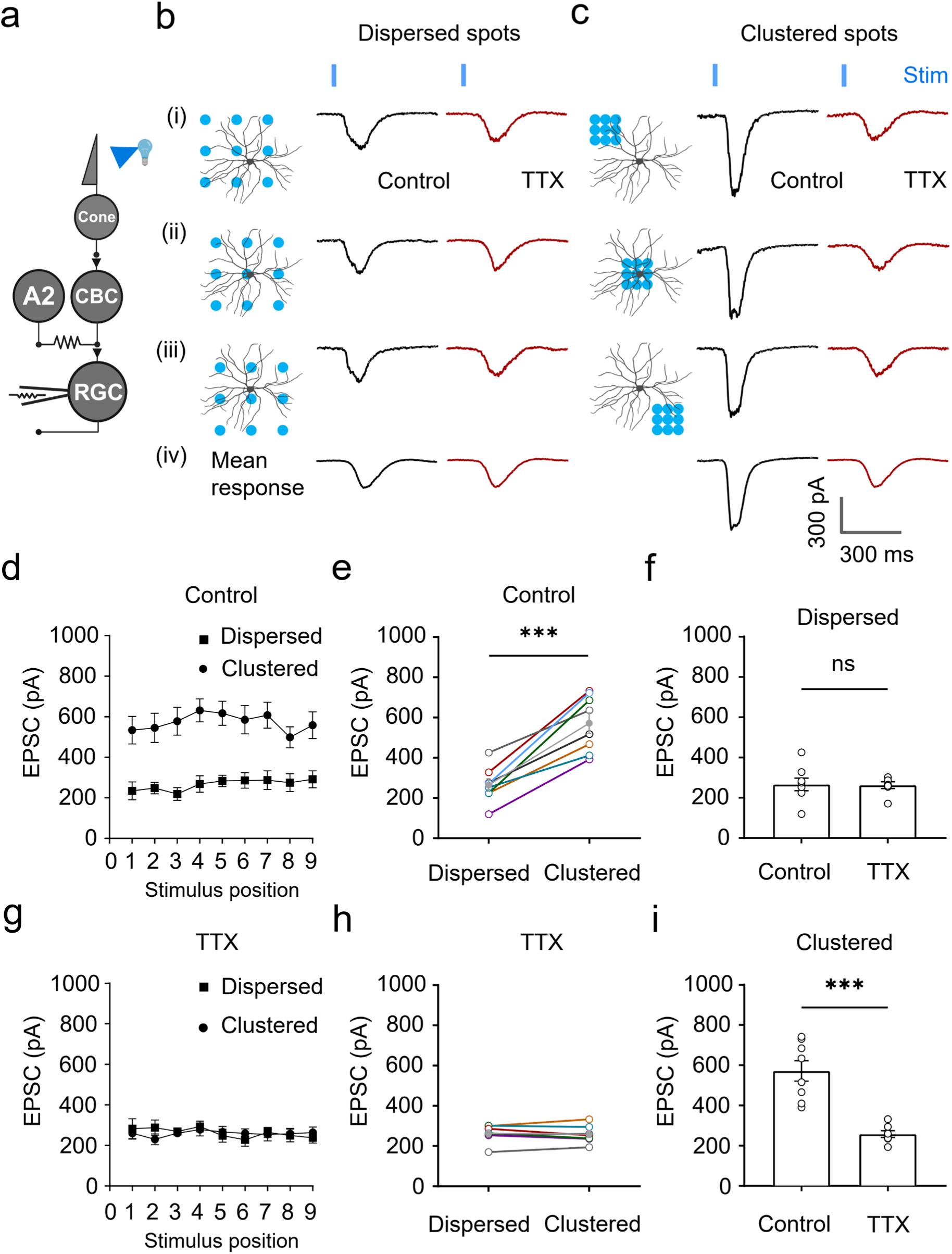
NaV-mediated excitability amplifies natural RGC light responses driven by spatially adjacent cones. **a)** Retinal circuit diagram, showing experimental arrangement. The light-adapted retina was exposed to spots of light to activate cones while EPSC responses from an RGC were recorded under voltage clamp. **b)** A set of 9 dispersed spots, each 36 µm in diameter and separated by 72 µm was presented to the retina, all confined within the dendritic tree territory of a patch-clamped RGC that had been filled with fluorescent dye. The set of dispersed spots were presented in nine different positions relative to the RGC dendritic tree and the responses to three of which (i-iii) are shown. Note that all 3 positions elicited similar EPSCs, as did the mean response from all positions (iv). All of the responses were unaffected by TTX. **c)** Responses to the same 9 spots of light, but clustered to be immediately adjacent to one another. The clustered spots were presented in nine different positions and the responses to three (i-iii) are shown. Note that all 3 positions elicited larger EPSCs which were all reduced by TTX. **d, g)** Responses to dispersed and clustered spots without and with TTX at each position (two-way ANOVA; stimulus type × stimulus location: control, F(4,56)=0.8276, n=8, p=0.5124; TTX, F(2,23)=0.5830, n=7, p=0.5568; stimulus type: control, F(1,14)=26.50, n=8, p=0.0001; TTX, F(1,12)=0.0194, n=7, p=0.8916). **e, h)** Average response at all 9 positions. Note that the clustered spots produced almost 2-fold larger EPSCs compared to the dispersed spots (clustered=573 ± 51 pA, dispersed=266 ± 31 pA, n=8 each; p=0.0002; paired t-test). In the presence of TTX, however, clustered spots elicited smaller EPSCs which were roughly the same size as produced by the dispersed spots (clustered=259 ± 16 pA, dispersed=262 ± 16 pA, n=7 each; p=0.7350; paired t-test). (**f, i)** Comparison of all responses to dispersed and clustered spots, without and with TTX (dispersed: control=266 ± 31 pA, n=8, TTX=262 ± 16 pA, n=7; p=0.9215; unpaired t-test; clustered: control=573 ± 51 pA, n=8, TTX=259 ± 16 pA, n=7; p=0.0003; unpaired t-test). Data in f, i are mean ± sem. These experiments were carried out in mesopic background (758 R*/rods/s) at 100 % contrast. The duration of stimulus was 10 ms.

We stimulated cones with an array of 9 spots of light, each ∼36 µM in diameter. The 9 spots were either widely dispersed, with each spot 72 µm from its nearest neighbor (left, Fig. 6b), or tightly clustered, with the 9 spots immediately adjacent to one another (left, Fig. 6c). In both cases, the array of spots was positioned to fall within an area delineated by the dendritic tree of the On-alpha RGCs. In principle, both dispersed and clustered stimulus arrays should activate the same number of pre-synaptic BCs, and with no extra amplification, should generate the same summated EPSC.

Not surprisingly, the dispersed array of spots generated a moderate-sized summated EPSC that was unaffected by adding TTX (Fig. 6b, f). However, we found that the clustered array generated a summated EPSC that was almost twice as large as predicted (dispersed=266 ± 31 pA; clustered=573 ± 51 pA; n=8 each; p=0.0002) and had a faster rise time (dispersed=108.2 ± 6.1 ms; clustered=87.4 ± 5.7 ms; n=8 each; p=0.0056) (Fig. 6c, e). TTX had a dramatic effect on the response to the clustered array, slowing its rise time and reducing its peak amplitude to match that elicited by the dispersed array (Fig. 6h, i).

Shifting the dispersed and clustered arrays to different positions within the dendritic tree collecting area of the RGC made little difference to the relative amplitudes of summated EPSCs that were elicited. The response to the clustered array was always about twice as large as the dispersed array, whether the cluster fell on BCs cells near the center or the periphery of the RGC’s dendritic tree (Fig. 6d, g). The simplest interpretation of these results is that activation of neighboring cones results in activation of spatially clustered BCs, which converge onto common A2 cells, resulting in NaV-mediated synaptic amplification of CBC6 and potentially other On-CBCs output, whereas activation of dispersed cones does not lead to amplification.

All of the above experiments were carried out in mesopic lighting conditions, near cone threshold**^43^** (758 R*/rods/s). If A2-mediated amplification is an important mechanism for object detection, one might expect weaker amplification in brighter light. To evaluate the effect of lighting, we repeated the dispersed vs. clustered spot experiment while varying the intensity of both the background (I_B_) and the spot (I_S_) to maintain a constant contrast (I_s_/I_B_ = 2.0) (Fig. 7a-e). In mesopic lighting (background= 758 R*/rods/s), dispersed spots generated a small EPSC (Fig. 7c, d), but clustered spots generated a much larger EPSC (peak amplitude 2.2-fold larger). In low photopic light (background= 3,790 R*/rods/s, clustered spots elicited EPSC that were only modestly larger than with dispersed light (1.2-fold larger).

**Figure 7.**
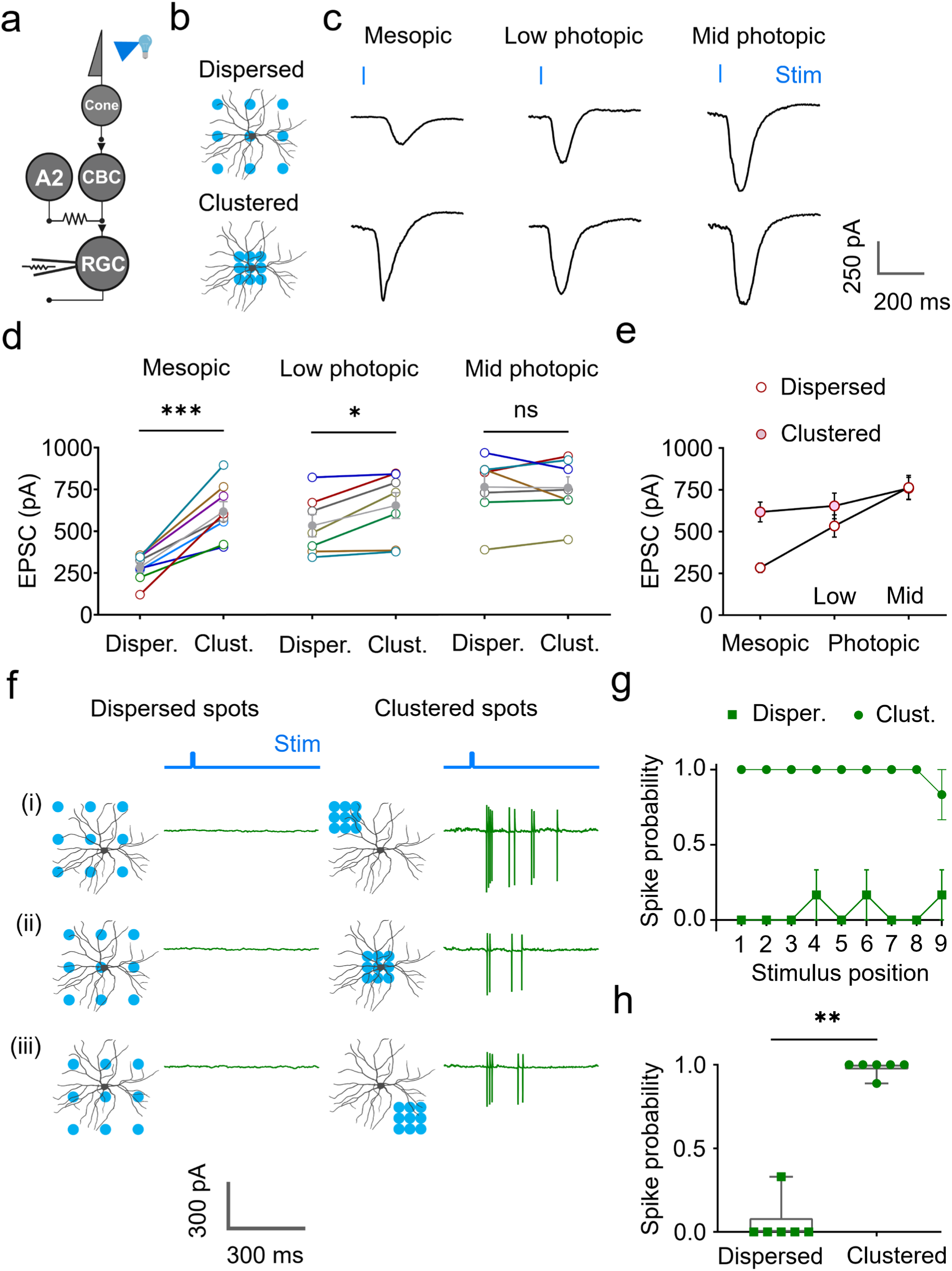
Amplification is largely restricted to mesopic conditions. **a)** Retinal circuit diagram, showing experimental arrangement. The retina was exposed to spots of light to activate cones while EPSC (c-e) or spike responses (f-h) from an On-alpha RGC were recorded. **b)** Dispersed or clustered spots were presented to the retina in one position **c)** EPSC recordings from an On-alpha RGC in response to the dispersed spots (top) and the clustered spots (bottom) with increasing background light: mesopic (758 R*/rods/s), low-photopic (3790 R*/rods/s) and mid-photopic conditions (18950 R*/rods/s). **d)** Clustering of spot stimuli strongly amplified responses in mesopic background (dispersed=283 ± 29 pA, clustered=617 ± 59 pA, n=8 each, p=0.0003, paired t-test), weakly in low-photopic background (dispersed=533 ± 67 pA, clustered=653 ± 77 pA, n=7 each, p=0.0169, paired t-test) and not at all in mid-photopic background (dispersed=763 ± 73 pA, clustered=759 ± 66 pA, n=7 each, p=0.9056, paired t-test). Open circles represent individual RGC responses; filled grey circles represent the mean response from all recorded RGCs. **e)** Background alters responses to the dispersed spots but not the clustered spots. EPSC amplitude to the dispersed spots increased from mesopic to low-photopic to mid-photopic background (One-way ANOVA; background light: F(2,19)=18.12, p=<0.0001). In response to the clustered spots, the size of EPSCs did not change across these backgrounds (One-way ANOVA; background light: F(2,19)=1.198, p=0.3236). Note that unlike the response to the dispersed spots, the response to the clustered spots was already nearly-maximal EPSCs in the mesopic background. For these experiments, contrast was 100 % in all three conditions and the duration of stimuli was 10 ms. **f)** Left: The dispersed spots were presented in nine different positions and the responses to three (i-iii) are shown. Note that none of the stimulus positions elicited spikes. Right: The clustered spots were presented in nine different positions and the responses to three (i-iii) are shown. Note that all 3 stimulus positions elicited spikes. **g)** Mean probability of spiking in response to the dispersed and the clustered spots at all 9 positions (two-way ANOVA; stimulus type × stimulus location: F(2,17)=0.8197, n=6 each, p=0.4384; stimulus type: F(1,10)=243.2, n=6 each, p˂0.0001). **h)** Cell-wise probability of spiking in responses to the dispersed and the clustered spots (dispersed=0.06 ± 0.06, clustered=0.98 ± 0.02; n=6 each; p=0.0022, Mann-Whitney test). For panels g and h, error bars denote ± sem. Note that these cell-attached spike recordings were performed in mesopic background at 100% contrast. Spiking probability denotes the fraction of stimulus that elicited spikes, which could vary between 0 to 1.

In brighter photopic light (background= 18,950 R*/rod/s), clustered spots generated EPSC responses with the same peak amplitude as dispersed spots (Fig. 7c, d). In general, RGC responses to dispersed spots gradually increased with increasing intensity, while the response to clustered spots remained largely unchanged (Fig. 7e). These results indicate that A2 cells strongly amplify responses in mesopic conditions, mildly in low photopic conditions. and not at all in mid-photopic or brighter conditions. The constancy of responses to clustered stimuli across several log units of light intensity is consistent with A2 cell-mediated amplification playing an important role in object recognition.

Finally, we asked whether the spatial clustering preference exhibited by A2 cells affects the spike output of RGCs, the critical signals for sensory processing by the brain and visual perception by the animal. We obtained “loose patch” extracellular recordings from On-alpha RGCs, and quantified the probability of spikes occurring in response to dispersed or clustered spots of light that activated cone photoreceptors. Once again, we found a strong preference for clustered stimuli, independent of the specific location within the center of the RGC’s receptive field (Fig 7f-h). This finding is consistent with the A2 amplifier playing an important role in tuning the response properties of On-alpha RGCs to respond more strongly to spatially distinct objects rather than spatially uncorrelated stimuli.

Up until now, our focus has been on the role of A2 amacrine cells in amplifying excitatory synaptic transmission between CBC6 cells and On-alpha RGCs. However, this is only one of many On-subcircuits in the inner retina. Several other types of On-cone bipolar cells are also electrically coupled to A2 amacrine cells **^21^** and along with CBC6 cells, provide inputs to most of the >20 On-RGCs in the mouse retina**^20,44^**. To test whether NaV channel-mediated spatial thresholding from A2 cells applies to other types of On-RGCs, we obtained patch clamp recordings from RGC somata that were located very close to presumed On-alpha RGC somata (within ˂50 µm). This makes these cells very unlikely to also be On-alpha RGCs, which are spaced at a regular inter-somatic distance in the mouse retina**^45^**. Dye-filling confirmed that the soma size and the dendritic tree pattern of these “non-alpha” On-RGCs were distinct from On-alpha RGCs (Fig 8 a). As we used Kcng4-cre X Ai9 mice for recording natural light responses, identification of alpha-RGCs, and therefore non-alpha RGCs, was more straightforward in this case as tdTomato expression was confined to alpha-RGCs**^46^**.

**Figure 8.**
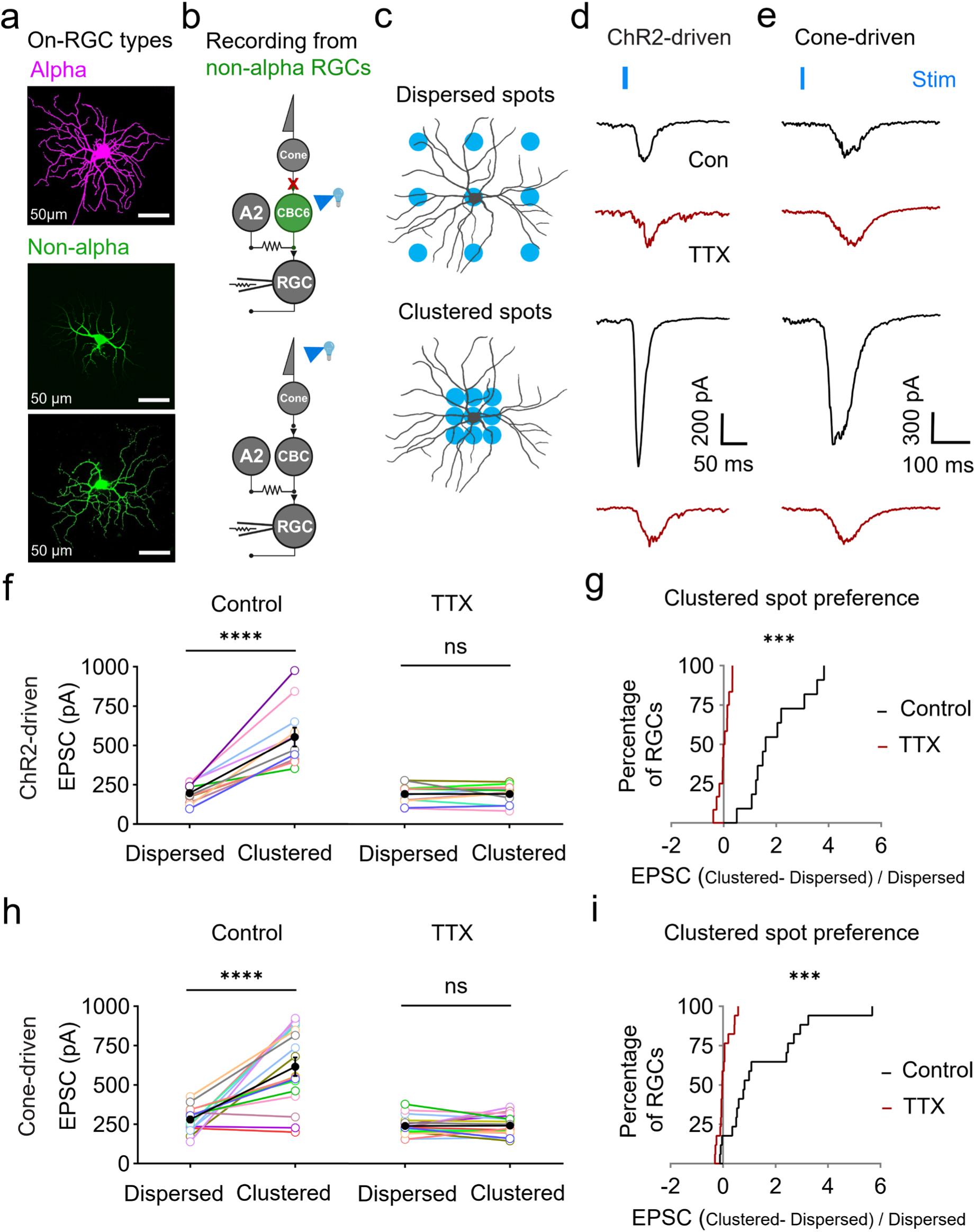
Multiple types of On-RGCs prefer spatially clustered stimuli, mediated by NaV-mediated excitability of A2 amacrine cells. **a)** Examples of dye-filled alpha and non-alpha On-RGCs. **b)** Retinal circuit diagram, showing experimental arrangement. The retina was photostimulated optogenetically, with ChR2 expressed in CBC6 cells and input from photoreceptors blocked (top), or naturally, by photostimulating cones (bottom). EPSC responses were recorded in non-alpha On-RGCs. **c)** Dispersed or clustered stimuli were presented as described in previous figures. **d, e)** Responses in non-alpha On-RGCs to optogenetic photostimulation of CBC6 cells (ChR2-driven) or natural photostimulation of cones (cone-driven). Example responses to dispersed (top) and clustered spots (bottom) are shown. **f, h**) Responses to dispersed vs. clustered spots in many individual non-alpha On-RGCs. With both optogenetic and natural stimulation, the clustered spots elicited larger EPSCs than did the dispersed spots (ChR2-driven: dispersed=196 ± 19 pA, clustered=553 ± 60 pA, n=11 each; p˂0.0001; paired t-test. Cone-driven: dispersed=280 ± 19 pA, clustered=615 ± 59 pA, n=17 each; p˂0.0001; paired t-test). For both stimulation protocols, TTX reduced EPSCs from clustered spots, eliminating the preference for clustering (ChR2-driven: dispersed=191 ± 17 pA, clustered=191 ± 17 pA, n=12 each; p=0.9876, paired t-test. Cone-driven: dispersed=239 ± 15 pA, clustered=242 ± 16 pA, n=17 each; p=0.8555, paired t-test). Filled black circles and error bars are mean ± sem. **g, i)** Summary results showing that the preference for clustering is eliminated by blocking NaV channels with TTX. **g)** For ChR2-driven responses without TTX, 11 out of 12 cells had a >50% preference for clustered stimulation (defined as >50% larger EPSCs), but with TTX, 0 out of 12 cells had a >50% preference for clustered stimulation. **i)** For cone-driven responses without TTX, 13 out of 17 cells had a >50% preference for clustered spots, but with TTX only one cell preferred clustering. Note that natural photo-stimuli were delivered in mesopic background with 100% contrast. The duration of stimuli was 10 ms in all experiments.

We recorded EPSCs in these non-alpha On-RGCs, elicited either by optogenetically photo-stimulating CBC6 cells or by naturally stimulating cone photoreceptors (Fig 8 b-c). We found that the majority of non-alpha RGCs responded preferentially to the clustered stimuli in both sets of experiments, and this preference was abolished by TTX (Fig 8 d-i). Together, these data indicate that NaV channel mediated amplification from A2 cells is a common feature of inner retinal circuitry, accentuating representation of spatial proximity in many types of On-RGCs in a previously unappreciated manner.

## DISCUSSION

### Amacrine cell-mediated amplification of bipolar cell synaptic transmission

A2 amacrine cells are the crucial hub in the retina where signals from rods and cones merge to influence the firing of RGCs. In mammalian retinas, about 20–50 rods converge upon each RBC, 20–25 RBCs converge upon each A2 amacrine cell, and 6-10 A2 cells, via their connection with On-CBC terminals, drive each On-alpha RGC. Overall, signals from >10000 rods converge upon a single ganglion cell**^17,47-49^**. The high degree of convergence increases the amplitude of rod-driven light responses, while non-linear properties of the rod synapse diminish responses triggered by uncorrelated events, increasing signal to noise**^50-52^**.

The cone pathway, in contrast, exhibits much less convergence. Each cone bipolar cell receives input from 2-6 cones**^39^**, and each RGC receives input from <400 cone bipolar cells**^20^**. In dim (mesopic) light, where cones are just beginning to respond and rods are nearing saturation**^43,53^**, bipolar cell light responses are small, hampering detection above noise. Our results indicate that NaV-mediated excitability of A2 amacrine cells, communicated to On-CBCs through gap junctions, amplifies chemical synaptic transmission from On-CBCs to On-RGCs, which should improve detection of cone-driven signals. With weak stimulation of CBC6 cells, A2 cell-mediated amplification is profound, increasing the amplitude of EPSCs by as much as 5-fold (Fig. 2h). In contrast, amplification of responses to stronger stimuli is more modest, increasing EPSCs by roughly 20% or not at all. This suggests that amplification is restricted largely to mesopic background, where it is needed to improve stimulus detection, in contrast to photopic light, where amplification is unnecessary. In scotopic light, where rods rather than cones are generating light responses, A2 amplification of RGC synaptic input is also absent**^17^**.

### The role of electrical excitability

A2 cells were previously found to slightly accelerate On-CBC to RGC synaptic transmission**^17^** but our findings reveal a previously unappreciated function of electrical excitability in CBC6 cells, amplifying spatially adjacent signals independent of the position within the RGC receptive field where they are displayed. This capability resembles the sophisticated response properties of complex cells in the primary visual cortex**^54^**, many synapses further along in the visual pathway. In cortical complex cells, shape detection agnostic to position emerges from the hierarchical nature of the visual pathway, involving many processing steps with each set of neurons converging on a smaller set in each successive steps**^55,56^**. But here, only 2 synapses into the visual system, similar feature detection properties emerge, which can be explained by the thresholding property of electrical excitability. This accentuates spatially correlated stimuli from neighboring presynaptic bipolar cells. Amplification only occurs if there are enough convergent inputs to bring the A2 cell to spike threshold, accounting for the preference for spatially clustered stimuli.

While our study focused on A2 amacrine cells, other narrow-field amacrine cell types, including the A8 amacrine cell, also has electrical synapses on type 6 bipolar cells**^57^**. Likewise, several other types of On-CBCs are also electrically coupled to A2 amacrine cells, including the type 5a and 7 On-CBCs**^21^**. Amongst the On-CBCs, type 5a, 6 and 7 bipolar cells constitute the majority of gap junctions on A2 cell**^21^**, and like most bipolar cells are devoid of NaV channels (Extended Data Fig. 5). Altogether, these cells comprise of almost 60% of On-CBCs**^20^**, stratify in three inner plexiform layers**^58^**, and likely provide excitatory input to a large fraction of On-RGCs. Consistent with this, our results show that NaV channel-mediated amacrine cell-driven amplification is not exclusive to On-alpha RGCs but applies to many other types of On-RGCs (Fig. 8). Hence the mechanism we have revealed is a common feature of inner retinal circuitry, accentuating spatial proximity in a previously unappreciated manner.

It has been suggested that the complex morphology of the A2 axodendritic tree endows the cell with distinct local and global integration capabilities**^59^**. However, the correspondence of the spatial extent of the axodendritic tree (∼36 um wide), and the spatial separation of visual stimuli that recruits NaV-mediated excitability (2 bars within ∼36 um; Fig. 5h), suggests that information integrated over the entire cell governs amplification of CBC6 cell output. Our findings hint at the function of the single “blind alley” appendage protruding from the soma of the A2 cells, which contains a high density of NaV channels, reminiscent of an axon initial segment**^60^**. In many brain neurons, spikes can be initiated independently in multiple dendritic branches, each comprising a computational unit that pre-processes synaptic inputs**^61^**. These dendritic spikes spread to the axon initial segment, which generates all-or none spikes that control neuronal output.

In A2 amacrine cells, the single central NaV channel-containing appendage suggests that the A2 cell comprises a single “computational unit”, at least under the experimental conditions used in this study. In this scenario, synaptic inputs that are spatially distributed across the dendritic tree of an A2 cell are integrated in one central location near the soma, which generates NaV-mediated spikes that back-propagate throughout the axodendritic tree. Presumably, these spikes reach chemical synaptic outputs onto Off-CBCs and electrical synapses onto On-CBCs, in both cases amplifying synaptic output.

### The role of gap junctions

We found that A2-mediated amplification depends critically on gap junctions that electrically couple CBC6 cells and A2 cells, which are located where fine processes of A2 cells intersect with synaptic terminals of CBC6 cells**^29^**. Pharmacological uncoupling of gap junctions with MFA had two dramatic effects on the physiology of CBC6 cells. First, uncoupling increased the input resistance of the CBC6 cell, such that small membrane currents resulted in much larger changes in membrane potential, as noted previously**^37,62^**. This leads to an increase in calcium-dependent neurotransmitter release, increasing the gain of synaptic transmission to RGCs (Fig. 2h). Second, uncoupling gap junctions removed NaV-mediated amplification of CBC6 synaptic transmission, decreasing the gain of CBC6 to RGC synaptic transmission. Interestingly, these two effects tend to offset one another, at least under our experimental conditions. Without the boost provided by the A2 amplifier, synaptic gain of the CBC6 output synapse would be low, shunting voltage changes in CBC6 terminals and reducing transmission. The rod pathway is highly convergent, such that many rods synapse onto far fewer rod bipolar cells (RBCs) and many RBCs synapse onto still fewer A2 amacrine cells, aiding in signal amplification. But the cone pathway exhibits much less convergence, making the A2 amplifier especially important for downstream signal detection.

Under natural conditions, the gap junctional coupling between A2 and On-CBCs is variable, subject to regulation by the neuromodulators dopamine and nitric oxide (NO) leading to uncoupling of gap juctions**^63,64^**. Dopamine and NO are released from distinct classes of amacrine cells whose activity is regulated by background illumination**^65,66^**. In bright background light or photopic conditions, dopamine and NO release is high, and gap junctions are uncoupled. This makes intuitive sense, as signaling through the cone pathway is strong in photopic light, making A2-mediated amplification unnecessary. In very dim background light or scotopic conditions, rods are active, but the rod pathway is not subject to A2-dependent amplification because A2 cells and RBCs are not electrically coupled. Under photopic conditions, A2-dependent amplification should be minimized by dopamine-dependent uncoupling of A2 cells from On-CBCs**^40^**. Hence, we expect that the A2 amplifier will have the greatest effect under mesopic conditions, when rod signaling is saturated but cones are barely exceeding threshold for responding to light**^43^**. It is precisely in this situation where amplification through the cone pathway should be most important for signal detection by downstream circuitry. Rod photoreceptors and rod bipolar cells are absent from primate fovea**^67^**, which may make amplification through this pathway particularly important for human mesopic vision.

Synaptic amplification by gap junctional-mediated sharing of excitability may be a common phenomenon. In the auditory brainstem, excitatory neurons can augment the electrical excitability of gap junction-coupled inhibitory interneurons**^68^**. Likewise, in the spinal cord, inhibitory interneurons can enhance synaptic output of electrically-coupled excitatory neurons**^69^**. Throughout the nervous system, gap junctions and chemical synapses co-exist and work in concert to influence the output of excitatory cells**^70-72^**. Gap junctional sharing of excitability among inhibitory interneurons promotes temporal coincidence detection**^73^**. Our findings suggest that sharing of excitability via gap junctions can also promote detection of spatial continuity to enhance object recognition by the visual system.

---

**Extended Data Figure 1.**
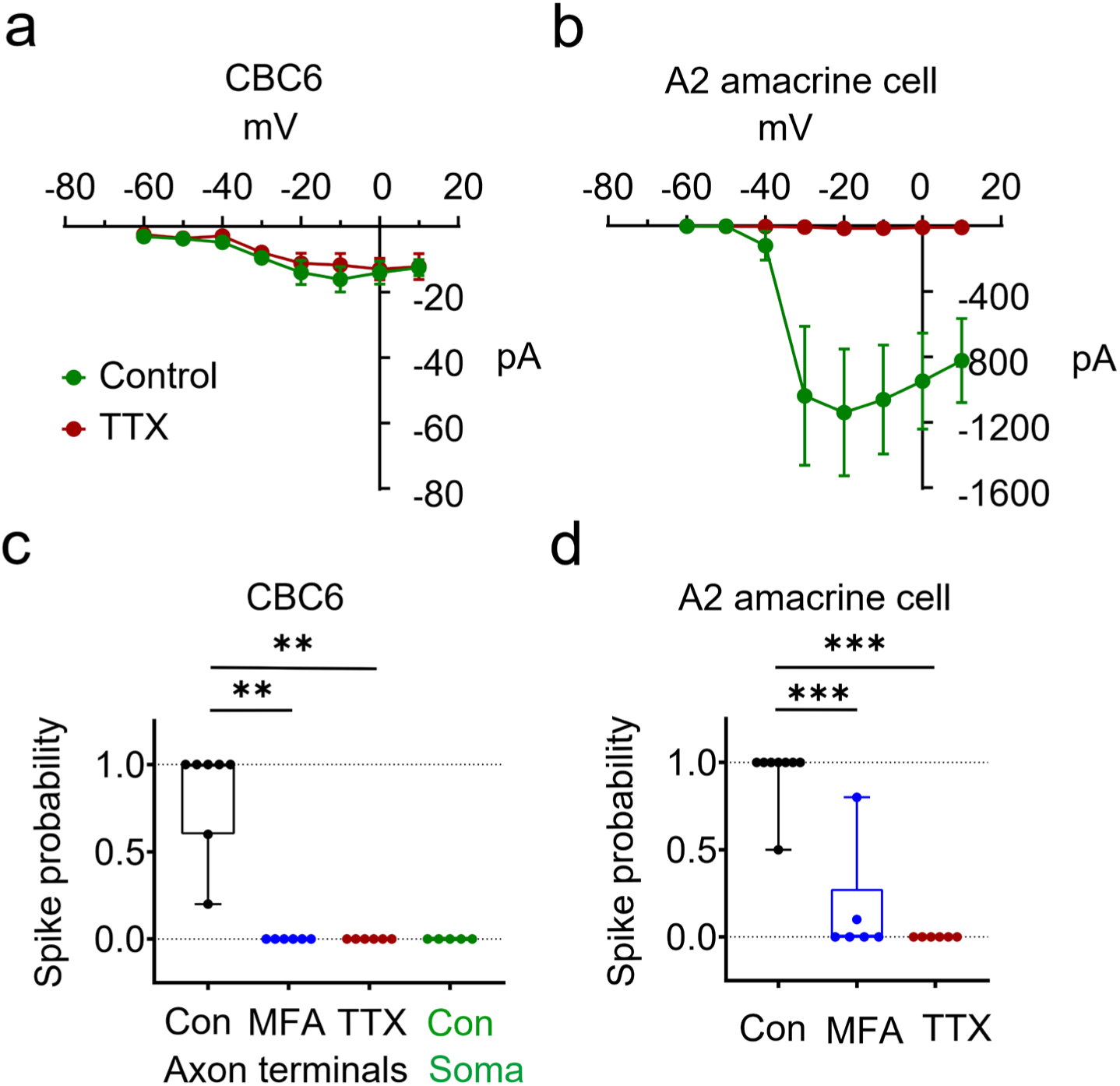
NaV-mediated excitability of A2 amacrine cells but not CBC6 cells. **a,b)** Current vs. voltage curves for peak inward current in CBC6s and A2 amacrine cells before and after addition of 1 µM TTX. At all voltage steps tested, TTX-inhibited NaV current in A2 cells was significantly larger than in CBC6 cells (two-way ANOVA: CBC6 cells, treatment × voltage steps, F(1,12)=0.1945, p=0.7559; treatment, F(1,8)=0.4910, p= 0.5034, n=5 each; A2 cells, treatment × voltage steps, F (1, 10)=7.495, p=0.0182; treatment, F(1,8)=8.729, p= 0.0183, n=5 each). **c)** Group data showing spike probability of CBC6 somata (0.00 ± 0.00, n=5) and CBC6 axon terminals in control, and after MFA or TTX (control=0.82 ± 0.11, n=7, MFA=0.00 ± 0.00, n=6, TTX=0.00 ± 0.00, n=6). Note that the probability of spiking in the CBC6 cell terminals was abolished by MFA or TTX application (Kruskal-Wallis test, p˂0.0001; control vs. MFA, p=0.0012; control vs. TTX, p=0.0012; Mann-Whitney test). **d)** Spike probability of A2 cells in control, and after MFA and TTX (control: 0.93 ± 0.06, n=8; MFA: 0.15 ± 0.13, n=6; TTX: 0.00 ± 0.00, n=6). Note that the probability of spiking in the A2 cells declined significantly after MFA treatment and was completely abolished by TTX (Kruskal-Wallis test, p˂0.0001; control vs. MFA, p=0.0007; control vs. TTX, p=0.0003; Mann-Whitney test). CBC6 cells expressed ChR2 and light flash intensity of 0.24 mW/cm^2^ and 10 ms duration were used to optogenetically stimulate these cells. Spike probability denotes the fraction of stimulus that elicited spikes, which could range from 0 to 1. Box plots depict median, min and max values. Data in a, b are mean ± sem.

**Extended Data Figure 2.**
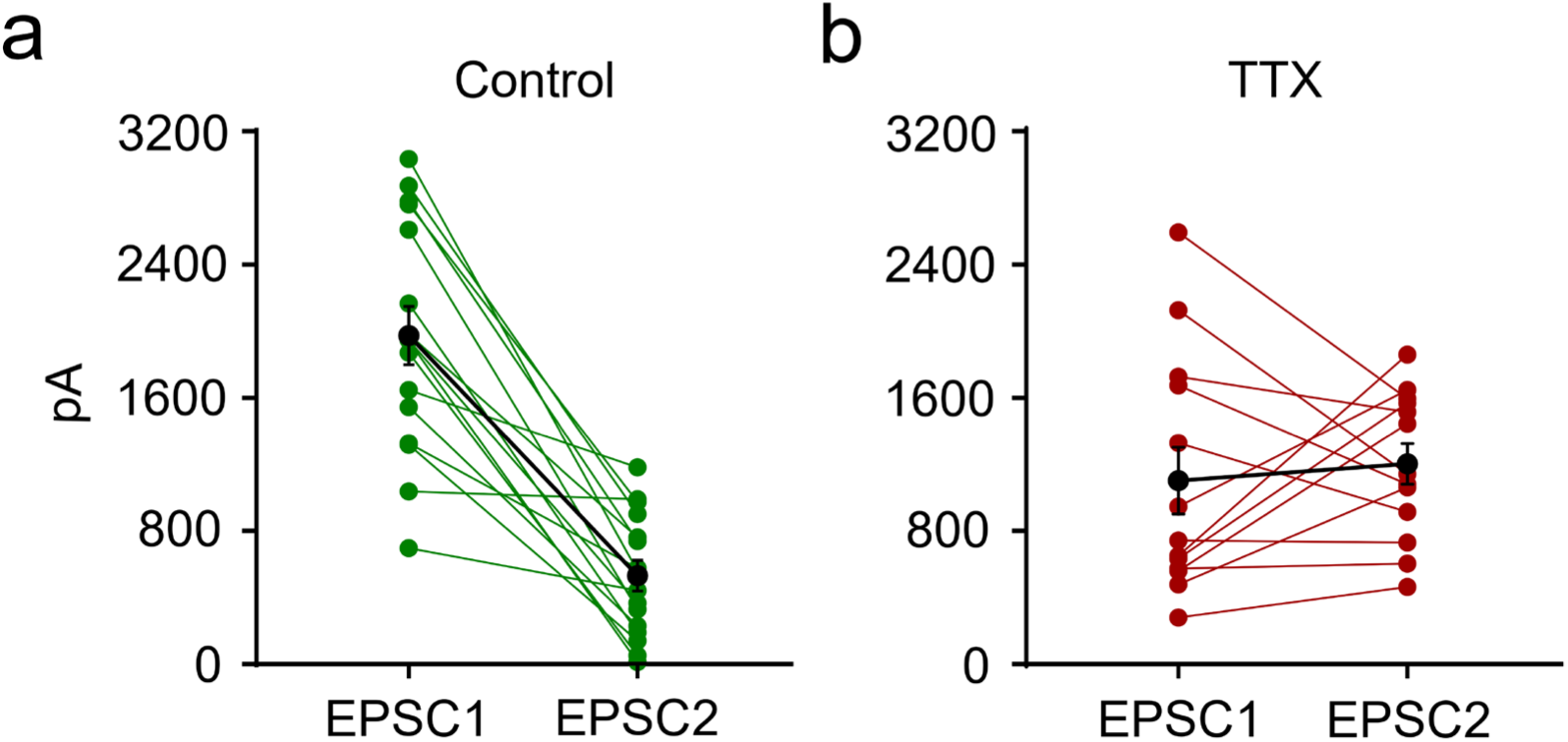
TTX changes short-term synaptic plasticity (paired-pulse behavior) from depressing to not depressing at the CBC6 cell to On-alpha RGC synapse. **a, b)** EPSCs elicited by two optogenetic stimuli (2ms light flashes) with an inter-stimulus interval of 50 ms before (a) and after (b) adding TTX (1 µM). Green and red points and lines represent individual recordings, black points and lines represent mean ± sem values. Note that paired EPSCs decremented without TTX, but grew in amplitude with TTX (control, n=16, TTX, n=13).

**Extended Data Figure 3.**
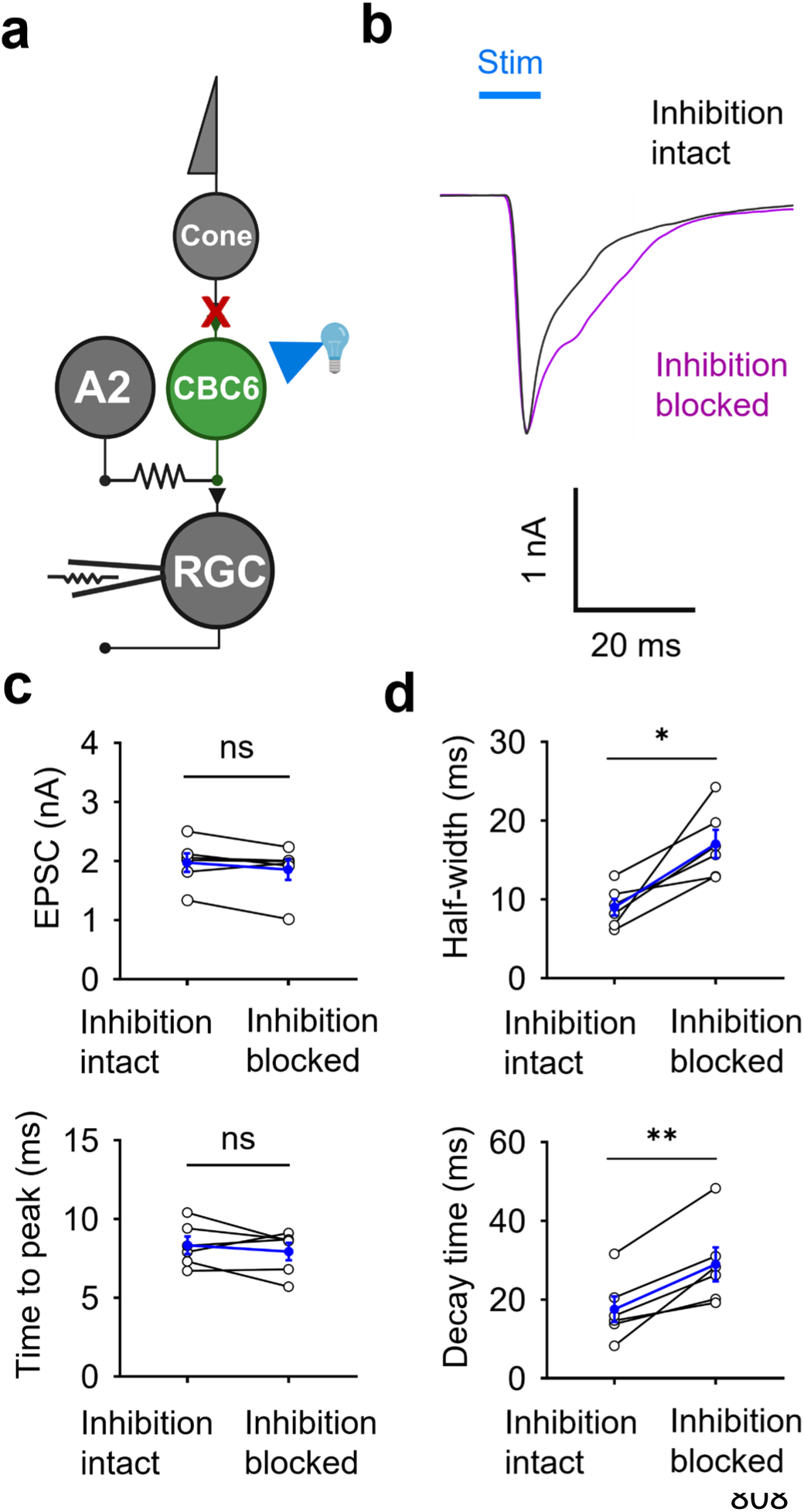
Inhibition does not alter the amplitude or time to peak EPSC. **a)** Retinal circuit diagram showing the experimental arrangement. Full-field light flashes at an intensity of 8.01 mW/cm^2^ were used to optogenetically stimulate CBC6 cells and whole-cell patch clamp recordings were obtained from On-alpha RGCs to measure evoked EPSCs under voltage-clamp. **b)** Example EPSCs recorded from On-alpha RGCs with inhibition intact (black) and blocked (magenta). Inhibition was blocked with antagonists of GABAa receptors (gabazine), GABAc receptors (TPMPA) and glycine receptors (strychnine). **c)** Inhibitory receptor blockers did not significantly alter the peak amplitude of EPSCs (inhibition intact=1.97 ± 0.15 nA, inhibition blocked=1.85 ± 0.17 nA, n=6 each; p=0.1562, Wilcoxon signed rank test) not did it affect time to peak (inhibition intact=8.33 ± 0.55 ms, inhibition blocked=7.93 ± 0.55 ms; p=0.4435, paired t-test). **d)** However, blocking inhibition prolonged EPSCs in every recording, both in terms of their half-width (inhibition intact=9.01 ± 1.05 ms, inhibition blocked=17.02 ± 1.78 ms; p=0.0122, paired t-test) and their 90%-20% decay time (inhibition intact=17.46 ± 3.25, inhibition blocked=28.86 ± 4.30 ms; p=0.0053, paired t-test). Blue circles and error bars indicate mean ± sem. Prolonged half-width and decay time suggests a delayed contribution of a non-A2 amacrine cell (presumably wf-Acs) in shaping the output from CBC6 cells.

**Extended Data Figure 4.**
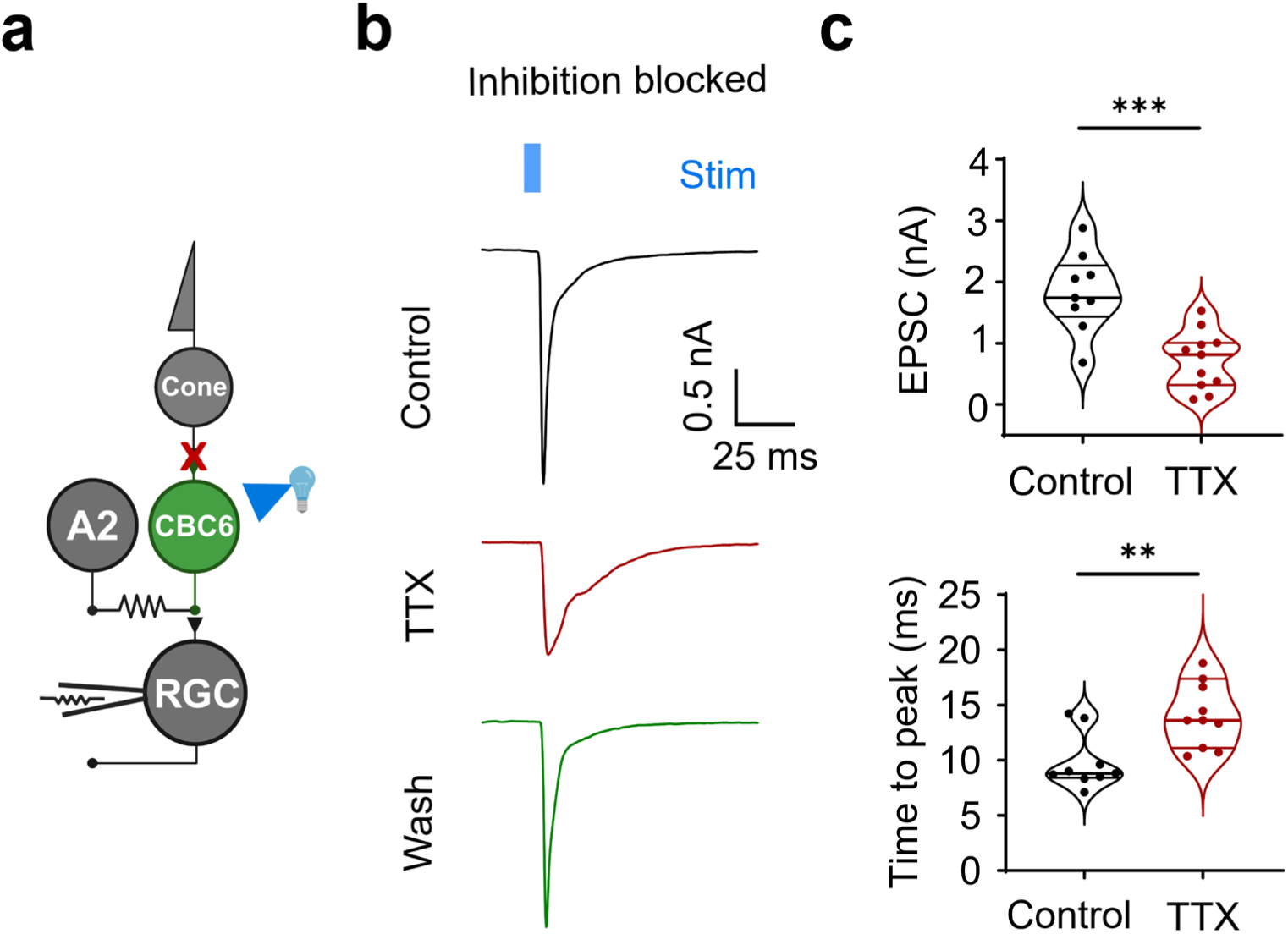
TTX reduces the amplitude and prolongs the time to peak of EPSCs even in the absence of inhibition. **a)** Retinal circuit diagram showing the experimental arrangement. Full-field light flashes at an intensity of 8.01 mW/cm^2^ were used to optogenetically stimulate CBC6 cells and whole-cell patch clamp recordings were obtained from On-alpha RGCs to measure evoked EPSCs under voltage clamp. **b)** Example EPSCs recorded from On-alpha RGCs in the absence (top and bottom) and the presence of 1 µM TTX (middle). Inhibition was blocked throughout these experiments with antagonists of GABAa (gabazine), GABAc (TPMPA) and glycine receptors (strychnine). **c)** Group data showing that EPSC amplitude and time to peak were altered by TTX, even though inhibition was blocked. TTX significantly reduced the peak amplitude (control=1828 ± 214 pA, n=9; TTX=721 pA ± 159 pA, n=11; p=0.0007; unpaired t-test) and delayed the time to peak (control=9.8 ± 0.8 ms; TTX=16.3 ± 2.4 ms; p=0.0029; Mann-Whitney test).

**Extended Data Figure 5.**
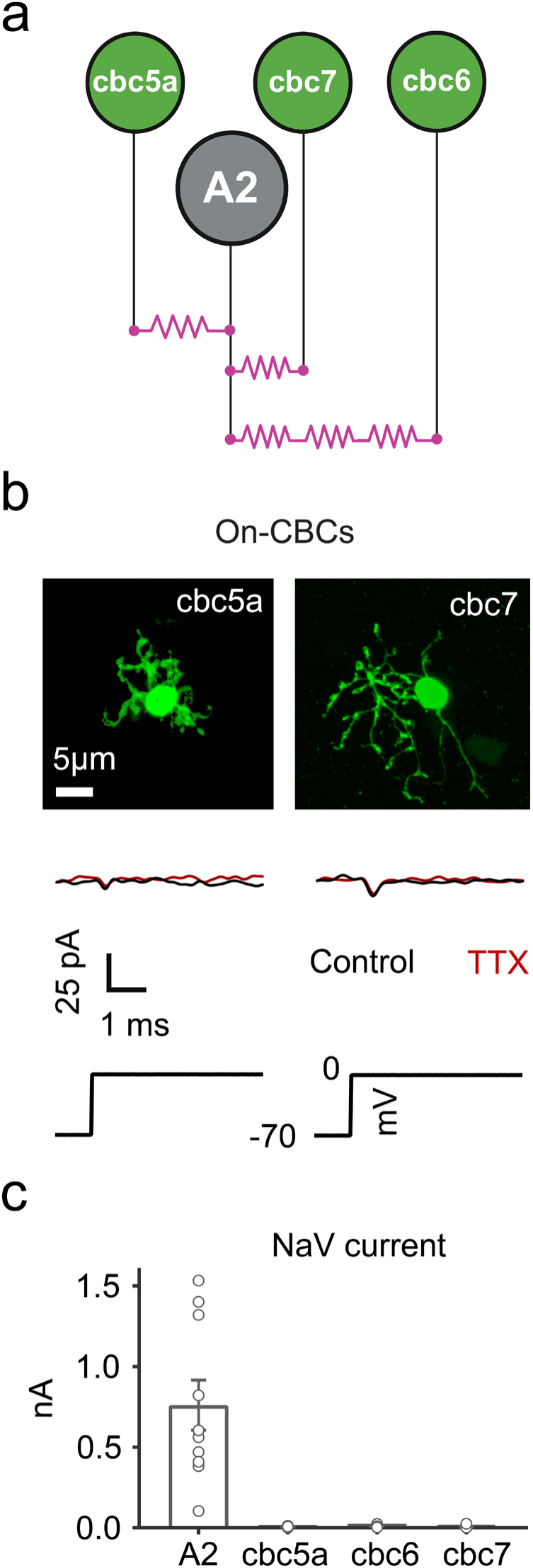
Other On-CBCs that are electrically coupled to A2 amacrine cells lack NaV channels. **a)** Schematic of gap junction circuitry between On-CBCs (green) and A2 amacrine cells. Gap junctions are highlighted in magenta. **b)** Examples of dye-filled On-CBCs in the Kcng4-cre X Ai9 mice; cbc5a (left), cbc7 (right). These cells were identified based on tdTomato expression**^19^**, axonal arbor size and stratification pattern**^74^**. Under voltage-clamp, cbc5a and cbc7 cells hardly exhibited depolarization-gated inward current (bottom), indicating lack of NaV channels just like their counterpart, cbc6 cells. Note that gap junctions, voltage-gated Ca^2+^ and K^+^ channels were blocked prior to NaV current recordings. **c)** Comparison of depolarization evoked inward or NaV current in A2 cell and its major gap junction partners (A2= 761 ± 155 pA, n=10; cbc5a= 5.2 ± 1.7 pA, n=8; cbc6= 11.3 ± 2.6 pA, n=8; cbc7= 6.4 ± 3.2 pA, n=7). Note that cbc5a and cbc7, together with cbc6 cells constitute almost 60% of On-CBCs**^20^** and 90% of bipolar cell gap junctions on A2 cell**^21^**, implying that multiple On-subcircuits are subject to A2-driven amplification as shown in Figure 8.

## Supplementary information

### Contribution of gap junctions to the passive membrane properties of CBC6 cells

By removing NaV-mediated amplification, one might expect that uncoupling CBC6 cells from A2 cells would reduce the amplitude of EPSCs recorded in the RGCs. However, EPSC amplitude either increased or remained unaltered after MFA application (Fig. 2, 3). For example, MFA increased the peak EPSC triggered by 1 ms optogenetic stimulation by almost 2-fold (control=965 ± 159 pA, n=16; MFA=1794 ± 104 pA; n=24; p=0.0002). A previous study showed that gap junctions account for a large fraction of the resting membrane conductance of CBCs**^37^**. We reasoned that by decreasing the resting conductance, gap junction uncouplers would allow membrane currents elicited by optogenetic stimulation to generate larger voltage changes. This would increase activation of voltage-gated Ca^2+^ channels and increase neurotransmitter release, perhaps enough to offset the loss of the A2 cell amplifier. We tested this idea by applying voltage steps to CBC6 cells and found that MFA treatment increased their input resistance (control=0.79 ± 0.06 GΩ, n=11; MFA=3.50 ± 0.44 GΩ; n=10; p=0.0002) (Supplementary Fig. 1a, b). Consistent with this, optogenetic stimuli generated a much larger depolarization of CBC6 cells after MFA treatment (control= 4.14 ± 0.69 mV; MFA=13.50 ± 2.36 mV; n=6 each; p=0.0094) (Supplementary Fig. 2). MFA also increased the input resistance of CBC5A cells (control=0.98 ± 0.09 GΩ, n=11; MFA=4.55 ± 0.69 GΩ; n=8; p=0.0012) (Supplementary Fig. 3), another On-bipolar cell type that forms gap junctions with A2 cells**^19^**. However, MFA had no effect on the input resistance of rod bipolar cells, which are devoid of gap junctions (control= 3.31 ± 0.38 GΩ; MFA=4.17 ± 0.71 GΩ; n=10 each; p= 0.3258) (Supplementary Fig. 1c, d).

Next, we tested the effect of gap junction uncoupling on synaptic transmission from CBC6 cells. To remove any contribution from NaV channel-mediated amplification, TTX was present throughout these experiments. Under these conditions, 1 ms optogenetic stimulation of CBC6 cells elicited small EPSCs in RGCs, but these grew much larger after treatment with MFA (without MFA=158 ± 39 pA, n=14; MFA=1789 ± 116 pA; n=11; p<0.0001) (Supplementary Fig. 1e-g). Another gap-junction uncoupler, 18β-glychyrhhetinic acid (18β-GRA) also increased EPSCs in RGCs (Supplementary Fig. 4). However, MFA had no effect on EPSCs elicited in A2 cells upon rod bipolar cell photostimulation (without MFA=427 ± 93 pA, n=11; MFA=434 ± 81 pA; n=10; p=0.8633) (Supplementary Fig. 1h-j).

Together, these experiments reveal that gap junctions set the resting membrane resistance of CBC6 cells and other electrically coupled On-CBCs, and that depolarization whether elicited by physiological or optogenetic stimuli, is reduced by this gap junctional shunt. Activation of NaV channels in the A2 cell helps offset this signal reduction, ensuring that depolarization of CBC6 terminals can effectively activate voltage-gated Ca^2+^ channels and trigger neurotransmitter release.

**Supplementary Figure 1.**
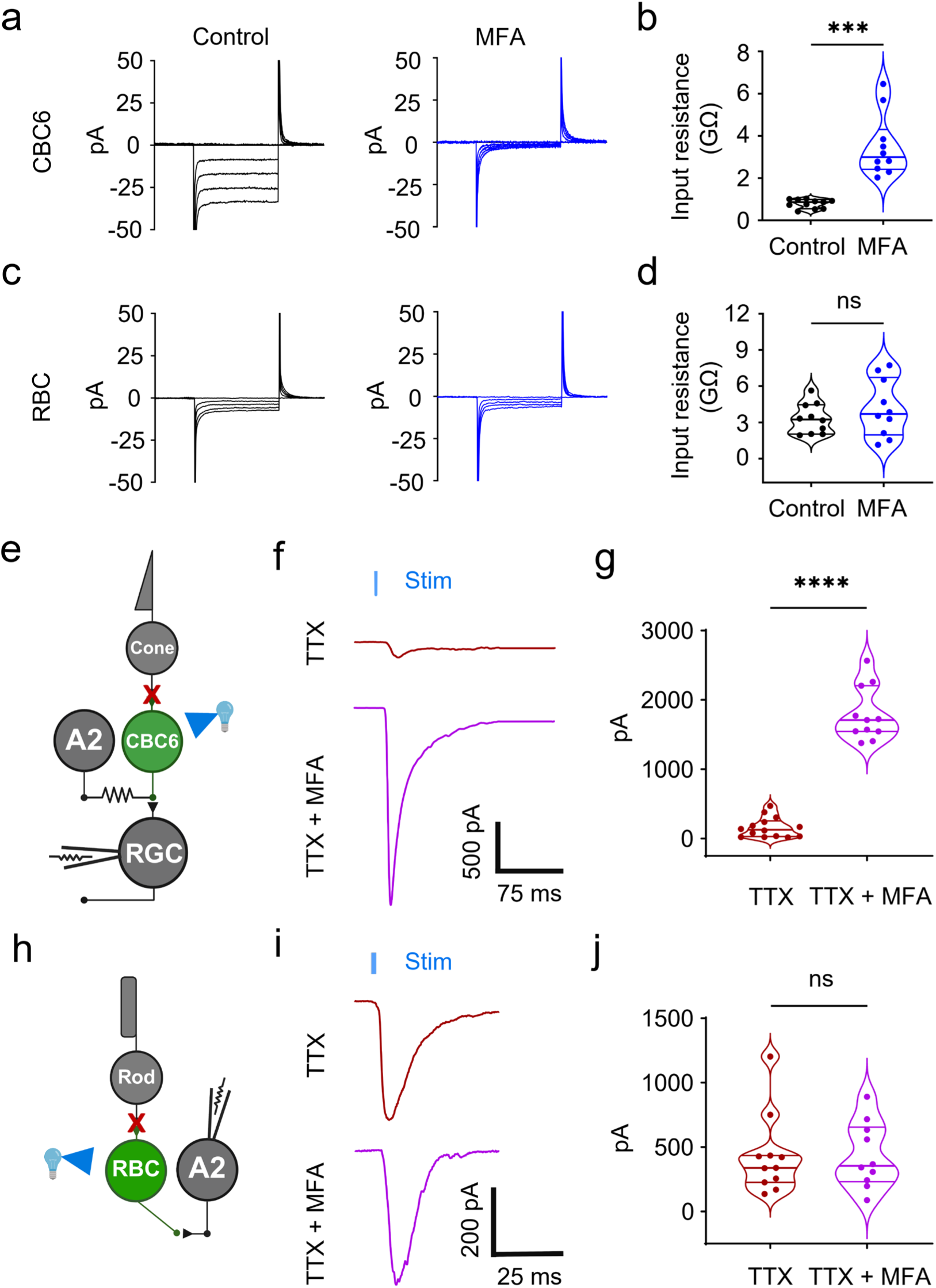
Gap junctions determine the input resistance of CBC6 cells. **a-d)** Membrane currents recorded from CBC6 cells (a) and RBCs (c) in response to a series of hyperpolarizing voltage steps (50 msec duration), with and without MFA pre-treatment to uncouple gap junctions. b, d) Steady-state input resistance (ΔV/ΔI), measured over the final 25 msec of the voltage steps. Note that MFA increased input resistance in CBC6 cells (b) but not in RBCs (d). MFA increased the input resistance of CBC6 cells by almost 4-fold (control=0.79 ± 0.06 GΩ, n=11; MFA=3.50 ± 0.44 GΩ; n=10; p=0.0002; unpaired t-test) but had no effect on the input resistance of RBCs (control=3.31 ± 0.38 GΩ; MFA=4.17 ± 0.71 GΩ; n=10 each; p= 0.3258; unpaired t-test). **e-j)** The gap junction shunt reduces synaptic efficacy of the CBC6 output synapse but not the RBC output synapse. **e, h)** Retinal circuit diagrams showing experimental arrangements. CBC6 cells or RBCs were optogenetically stimulated, eliciting EPSCs that were measured in the post-synaptic RGC or A2 cell. In all cases, TTX was present to block the NaV-dependent A2 cell amplifier. EPSCs elicited by stimulating CBC6 cells were enhanced by uncoupling gap junctions with MFA **(f, g)**, whereas EPSCs elicited by stimulating RBCs were unaffected **(h, i).** MFA enhanced CBC6 output by almost 11-fold (before MFA=158 ± 39 pA, n=14; after MFA=1789 ± 116 pA; n=11; p<0.0001; unpaired t-test) but had no effect on RBC output (before MFA=427 ± 93 pA, n=11; MFA=434 ± 81 pA; n=10; p=0.8633; Mann-Whitney test).

**Supplementary Figure 2.**
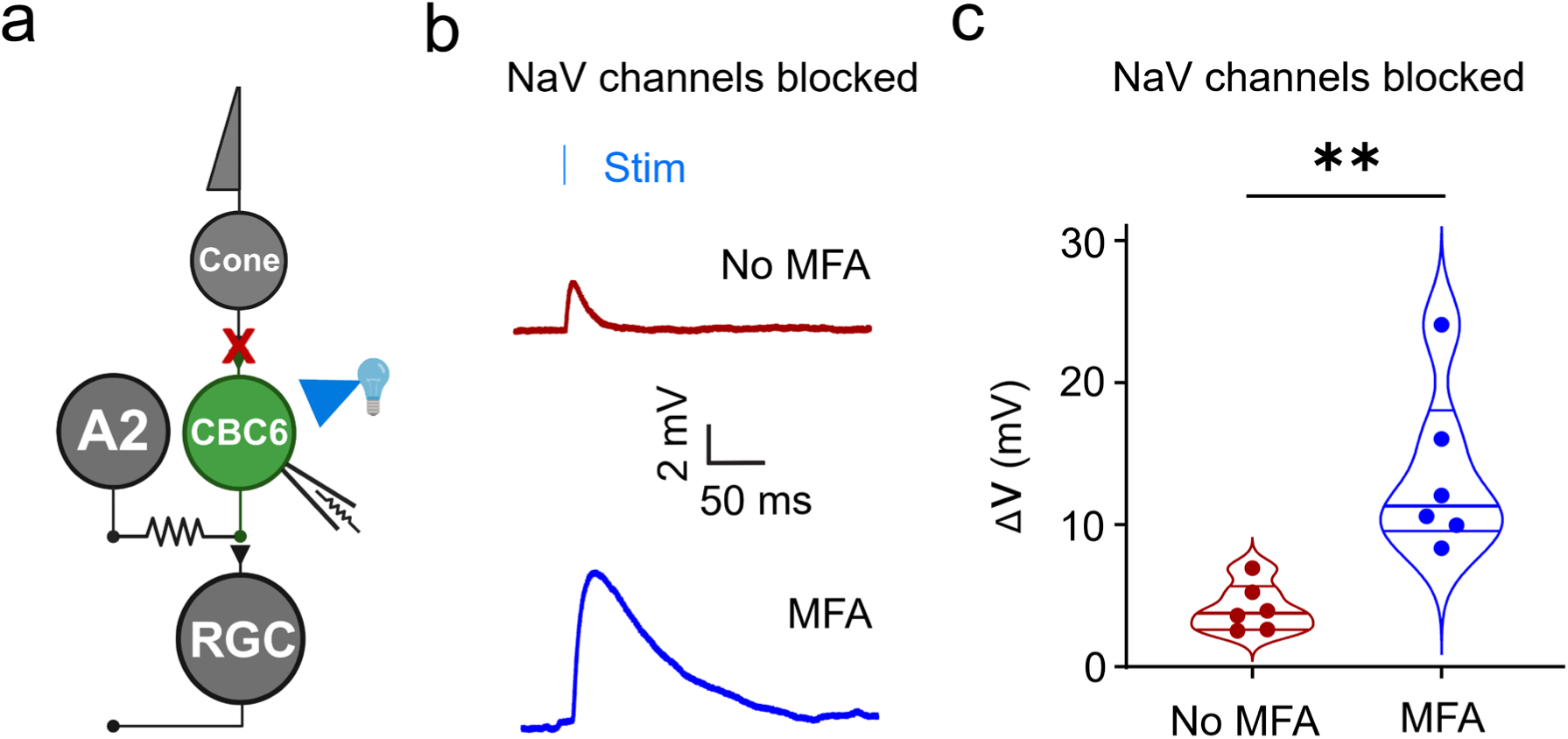
Uncoupling gap junctions enhances optogenetically-evoked voltage responses in CBC6s. **a)** Retinal circuit diagram showing the experimental arrangement. 1 ms full-field light flashes at an intensity of 1.21 mW/cm^2^ were used to optogenetically stimulate CBC6 cells and whole-cell current clamp recordings were obtained from a CBC6 cell to measure changes in membrane potential. **b)** An example recording from CBC6 cell before (top) and after treatment with MFA (bottom). NaV channels were blocked with TTX prior to recordings. **c)** Group data (violin plot) showing that MFA increases the evoked depolarizations by almost 3-fold (before MFA=4.14 ± 0.68 mV; after MFA=13.50 ± 2.36 mV, n=6 each, p=0.0094, unpaired t-test).

**Supplementary Figure 3.**
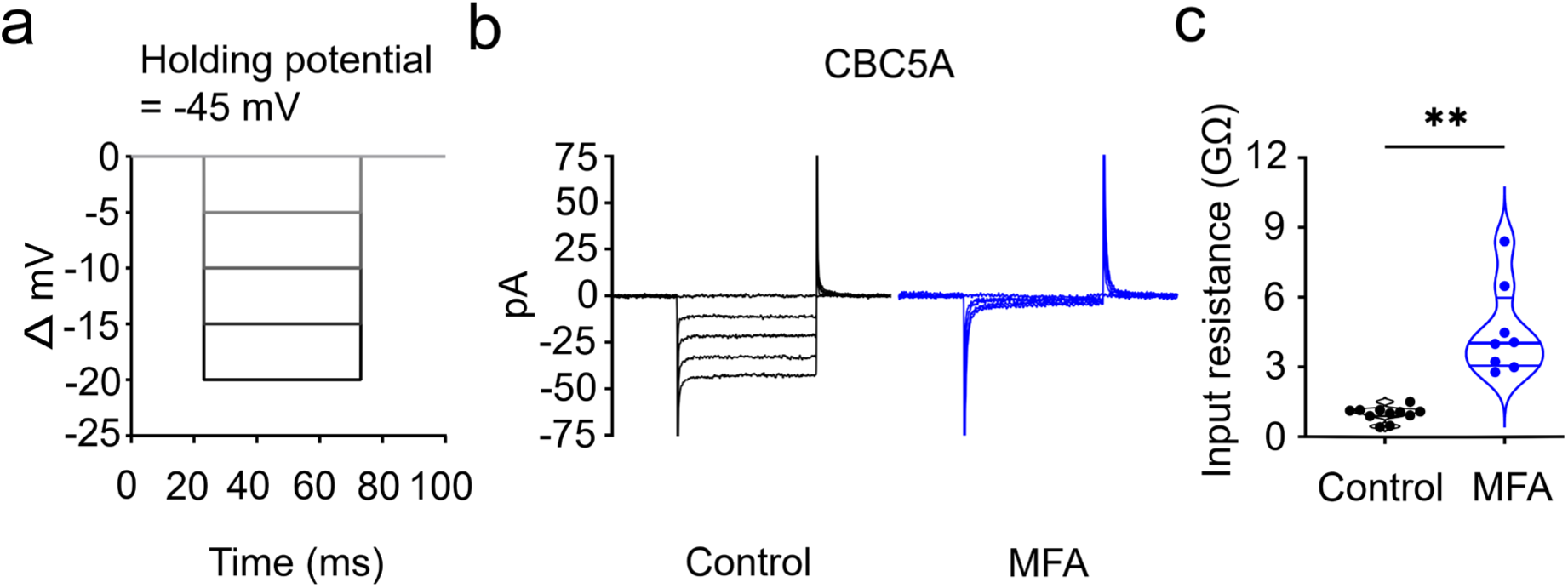
Gap junctions determine the input resistance of CBC5A cells. **a)** Input resistance was measured by delivering a series of hyperpolarizing voltage clamp pulses from -45 mV to -65 mV. **b)** Current responses in CBC5A cells before and after MFA treatment. **c)** Group data (violin plots) showing calculated steady-state input resistance. Note that MFA increased the input resistance of CBC5A cells by almost 4-fold (control=0.98 ± 0.09 GΩ, n=11; MFA=4.55 ± 0.69 GΩ; n=8; p=0.0012; unpaired t-test).

**Supplementary Figure 4.**
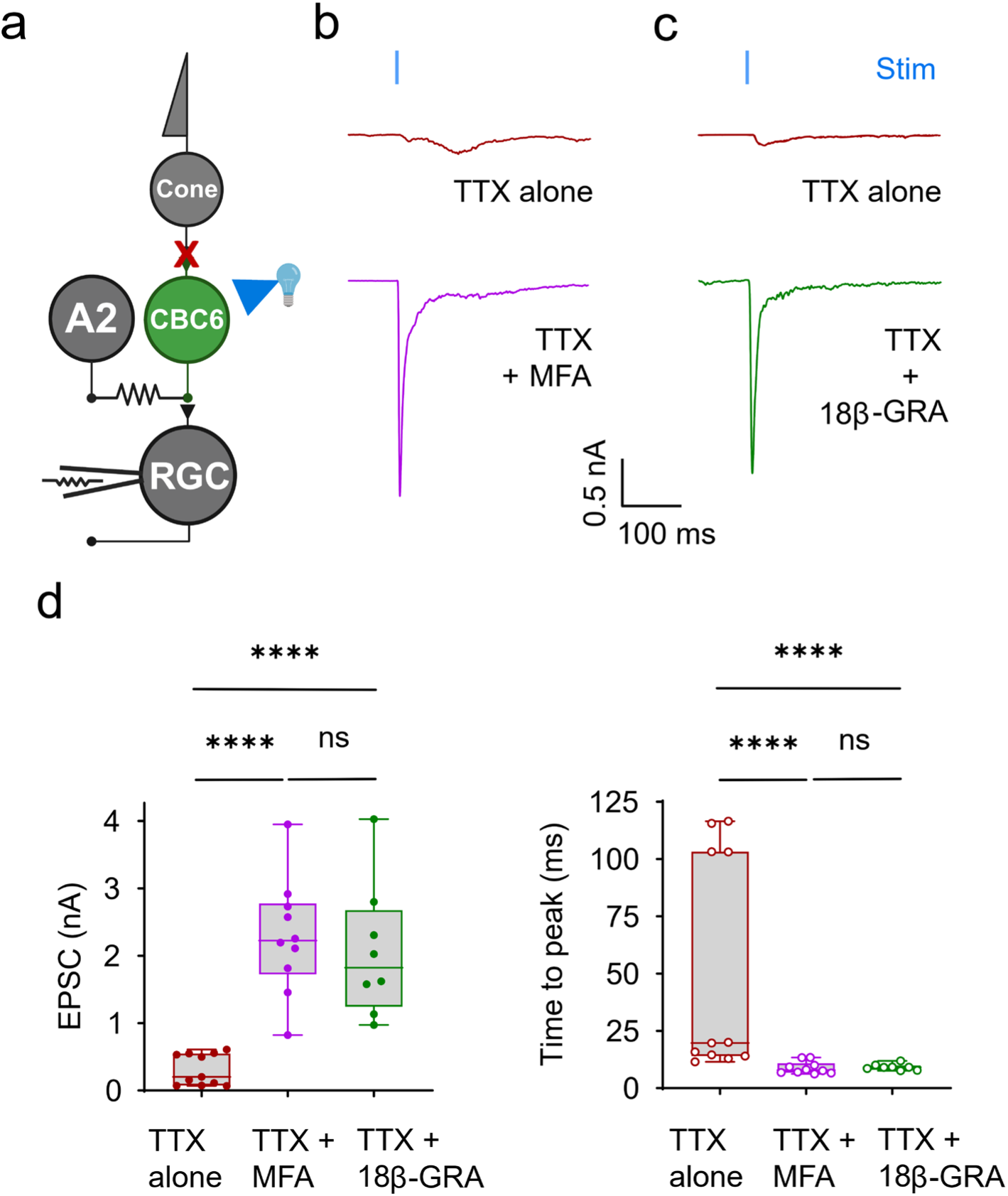
Uncoupling gap junctions increases evoked synaptic transmission at the CBC6 to On-alpha RGC synapse without NaV-mediated amplification. **a)** Retinal circuit diagram showing the experimental arrangement. Full-field light flashes (0.3ms) at an intensity of 8.01 mW/cm^2^ were used to optogenetically stimulate CBC6 cells and whole-cell patch clamp recordings were obtained from an RGCs to measure evoked EPSCs under voltage clamp. **b-c)** EPSCs elicited by a single flash were increased in amplitude after uncoupling gap junctions with MFA (**b**) or 18β-GRA (**c**). TTX was present throughout to block NaV-mediated excitability and eliminate A2 cell-mediated amplification. **d)** Group data (box plots) showing peak EPSC amplitude before (TTX alone, n=11), and 45 min after treatment with MFA (n=10) or 18β-GRA (n=8). Note that treatment with either MFA or 18β-GRA dramatically increased peak EPSC amplitude (Kruskal-Wallis test: p˂0.0001; post-hoc Mann-Whitney test: TTX alone=311 ± 70 pA, MFA=2282 ± 270 pA, 18β-GRA=2057 ± 352 pA; TTX alone vs. MFA, p<0.0001; TTX alone vs. 18β-GRA, p<0.0001) and shortened the time to peak (Kruskal-Wallis test: p=0.0001; post-hoc Mann-Whitney test: TTX alone=49.7 ± 14.4 ms, MFA=8.8 ± 0.8 ms, 18β-GRA=9.2 ± 0.5 ms; TTX alone vs. MFA, p<0.0001; TTX alone vs. 18β-GRA, p<0.0001).

## Methods

### Animals

Mice were handled in accordance with protocols approved by the UC Berkeley Institutional Animal Care and Use Committee (AUP-2016-04-8700-1) and conformed to the NIH Guide for the Care and Use of Laboratory Animals. CCK-ires-Cre knock-in mice or CCK-Cre mice (Jackson 012706) were used to drive Cre recombinase under the cholecystokinin promoter to label CBC6 cells. BAC-Pcp2-IRES-Cre or Pcp2-cre mice (Jackson 012706) were used to drive Cre recombinase under the control of mouse purkinje cell protein promoter to label RBC cells. Ai32 mice (Jackson 024109) were used to drive channelrhodopsin-2 or eYFP fusion protein in Cre-expressing CBC6 or RBC cells. The CCK-Cre, Pcp2-cre and Ai32 alleles were used in the mice in a hemizygous state. Likewise, Kcng4^-^Cre mice (Jackson: 029414) were crossed with Ai9 mice (Jackson: 007909) to drive tdTomato expression in type 5 bipolar cells and alpha RGCs. C57Bl/6J and Kcng4-Cre X Ai9 mice were used to record natural light responses from RGCs. Mice of both sexes were used interchangeably. 2 to 6 months old mice were used for all the experiments.

### Retinal Dissection

Mice were euthanized with isoflurane exposure and cervical dislocation. Eyes were enucleated and the retinal dissection was performed in the ACSF (in mM: 119 NaCl, 26.2 NaHCO_3_, 11 Dextrose, 2.5 KCl mM, 1 K_2_HPO_4_, 1.3 MgCl_2_* 6H_2_O, 2.5 CaCl_2,_ 293 mOsm/Kg). ACSF was oxygenated (5% CO_2_, 95% O_2_) whenever used. The dissection was performed in room light conditions as described previously**^76,77^**. In order to record natural light responses from RGCs, the mice were dark adapted overnight. However, dissecting in photopic conditions presumably bleached photoreceptors. Therefore, RGC recordings were performed after a recovery time of 3 hours. For retinal whole-mount experiments, retinas were flattened with 4 relieving cuts. For retinal slice experiments, flattened retinas were mounted on 13 mm diameter 0.45 µm filter paper discs (MF-Millipore) and sliced to a thickness of either 200 or 300 µm with a Stoelting Tissue Slicer. Retinal slices were rotated 90 degrees and mounted within a vacuum grease well.

### Electrophysiology

The majority of experiments were performed in the whole-mount retinas. For these experiments, the isolated retinas were incubated in the ACSF containing hyaluronidase (9,800 U/mL) and collagenase (2,500 U/mL) for 15 minutes to facilitate penetration of the inner limiting membrane**^78^**. Retinas were mounted ganglion cell side up in a recording chamber and held in place with a harp (Warner Instruments). The retinas were continuously superfused with oxygenated (5% CO_2_, 95% O_2_) ACSF at a flow rate of 4 ml/minute at 33-34 degree Celsius. Reagents for ACSF were purchased from Thermo Fisher Scientific. The cells of interest were viewed under DIC optics or epifluorescence with an upright microscope (Olympus). A2 amacrine cells were identified with the help of DIC optics and/or mCherry expression. RBC and CBC6 cells were targeted with the help of eYFP expression. CBC5A cells were identified with the help of tdTomato expression. Alpha and non-alpha On-RGCs were targeted under DIC optics for optogenetic experiments. For recording natural light responses, On-alpha RGCs were identified with the help of tdTomato expression**^46^**.

NaV-currents (Fig. 1) and EPSC recordings (Supplementary Fig. 1) from A2 cells and paired cell experiments (Fig. 3) were conducted in retinal slices. NaV currents and optogenetically evoked-EPSCs from A2 cells were recorded in 300 µm thick slices. For the paired cell experiments, slightly thinner 200 µm retinal slices were used. Superficial layers were avoided and deeply-seated somata (>75 µm) were targeted for patch-clamp recordings or sharp-electrode injections. For A2 cell recordings, the slices were superfused at a rate of 4 ml/ minute at 33-34 degree Celsius, same as for the whole-mount retinas. However, for the paired cell experiments from A2 cells and RGCs (Fig. 3), the ACSF flow rate was 6 ml/ minute, which was maintained at 30 degrees Celsius. A slightly faster flow rate and lower temperature somehow enhanced the viability of On-alpha RGCs in the slices.

Electrodes were made with borosilicate capillary glass tubing with OD= 1.5 mm, ID= 1.17 mm (Warner G150TF-4). For RGC recordings, glass was pulled to a resistance of 3-5 MΩs using a Narishige pipette puller. For A2 and BC cell recordings, electrodes used typically had a resistance of 6-8 MΩs and 9-12 MΩs, respectively. For QX-314 injections into A2 cells, sharp electrodes with resistance of 40-50 MΩs were used. These pipettes or sharp electrodes were pulled with Sutter Instruments P1000.

For voltage clamp experiments, the pipette solution contained (in mM): 123 Cs gluconate, 8 NaCl, 1 CaCl_2_, 10 EGTA, 10 HEPES, 10 glucose, 5 Mg^2+^ ATP, 5 QX 314, 0.01 Alexa 594, pH 7.4 with CsOH, 293 mOsm/Kg. For current clamp experiments, the pipette solution contained (in mM): 125 K^+^ Gluconate, 10 EGTA, 10 KCl, 10 HEPES, 4 Mg^2+^ ATP, 0.01 Alexa 488 or 594, 293 mOsm/Kg. Alexa Fluor 488, 568 or 594 Hydrazide (ThermoFisher Scientific) at 10 µM was used to dye fill cells during recording for confirmation of cell identity. In some cases, 0.2% Neurobiotin was included in the recording pipette to confirm the identity of the cells post-hoc with immunostainings. Whole-cell recordings were sampled at 10 or 20 kHz and filtered at 2 kHz with a Multiclamp 700B amplifier, digitized with a 1440a or 1550a Digidata A-D converter and analyzed offline with Clampfit. Recordings with series resistance with values higher than 10% of the cell membrane resistance or 30 mΩ were discarded.

### Pharmacology or drugs used

In all the optogenetic experiments, the kainate receptor antagonist ACET (1 µM) and the mGluR6 agonist L-AP4 (10 µM) were added to the ACSF in order to block photoreceptor driven light responses. In addition, NMDA and acetylcholine receptors were blocked with D-AP5 (50 µM) and Curare (25 µM) respectively. Gabazine (10 µM), TPMPA (50 µM) and strychnine (10 µM) were added to the ACSF to block ionotropic GABA_A_, GABAc and glycine receptors when necessary (Extended Data Fig. 3, 4). Mibefradil (10 µM) and Nimodipine (10 µM) were used to block voltage-gated calcium channels (Fig. 1, Extended Data Fig. 1, 5). MFA (100 µM) and 18β-GRA (50 µM) was used to uncouple gap junctions. TTX (1 µM) was used to block voltage-gated sodium channels. In addition, QX-314 (5mM) was always included in the pipette while recording EPSCs from the RGCs. QX-314 (1mM) was injected into the A2 cells with the help of sharp electrodes (Fig. 3). Pharmacological agents were purchased from Tocris Biosciences.

### Light stimulation or projection

Optogenetic stimulation of CBC6s and RBCs was provided by a 470 nm LED delivering a maximum of 2.1X10^8^ photons/s, (Lumencore or CoolLED pE-4000) measured at the plane of the retina with a spectrophotometer (Thor Labs). When necessary, stimulation intensity was varied by changing the duration of the stimulus ranging from 0.1 to 10 ms. Thus, the number of photons delivered during the optogenetic stimulus ranged from 2.1 × 10^5^ (0.1ms or 1%) to 2.1 × 10^7^ (10ms or 100%). For optogenetic experiments in Fig. 4-5, 10 ms stimulus delivered 1 × 10^5^ photons/s.

For experiments in Fig. 4-8, bars or spots of light were generated with a Texas Instruments Digital Light Projector (CEL1015 Light Engine, Digital Light Innovations) controlled with CELconductor Control Software. Projection of bars or spots of light and simultaneous EPSCs or spike recordings were controlled by Digidata. All stimuli were provided through 40X or 60X Olympus objectives except when measuring natural light responses. In this case, 20X Olympus objective was used to map the entire dendritic tree of On-alpha RGCs with nine sets of dispersed or clustered spots. For these experiments, three different stimulus intensities were used: 3.92 × 10^3^, 1.98 × 10^4^ and 9.92 × 10^4^ photons/µm^2^/s. The respective background light intensities were: 1.96 × 10^3^, 9.92 × 10^3^ and 4.96 × 10^4^ photons/µm^2^/s. Light intensities were converted into R*/rods/s as described previously**^79^**. In each case, the contrast was 100%, which was calculated as follows: Contrast=(Stimulus intensity-Background intensity)/Background intensity.

### Viral injections

AAV-PHP.eB-HKamac-mCherry-T2A-TRPV1.PHP.e (10^13^ viral genome particles/mL) was produced in the Gene Delivery module of the Vision Science Core at UC Berkeley. Ocular injection of this construct resulted into expression of mCherry specifically in A2 amacrine cells as HKamac is a A2 cell-specific promoter**^80^**. We injected this viral construct into the eyes of Cck-Cre-Ai32 mice in order to target A2 cells. In short, mice were anesthetized with isoflurane. A small hole was made at the margin of the cornea and sclera by using a 30-gauge needle. Approximately, 1.5 μL of the virus was intravitreally injected into the eyes of the mice. After injection, mice were returned to their home cage and monitored, until fully recovered. Retinas were used for experiments three to four weeks after the injections.

### Immunohistochemistry and scanning

The dye and/or Neurobiotin injected retinas were fixed with 4% PFA. After washing steps, they were incubated with Alexa488 or 647 conjugated Streptavidin (S32354 or S32357, Thermo Fisher Scientific) as described before**^81^**. Images were scanned with 40X/1.4 oil objective using Zeiss LSM780 Laser Scanning Confocal Microscope. The z-stacks were acquired at a pixel size of 60 nm × 60 nm at a z-step of 0.3 µm or 1 µm. In some cases, images captured were deconvolved with Huygens Essential Software (Scientific Volume Imaging, Netherlands) as described before**^82^**.

### Data Visualization and Statistics

All reported statistics and figures were generated with GraphPad Prism9. Retinal and experimental diagrams were created with Biorender.com. Data are presented as mean ± sem; median and 25^th^ and 75^th^ quartiles; median, minimum and maximum. Normality was checked with Shapiro-Wilk test prior to applying statistical tests. Significance of non-normal data was tested with Mann-Whitney U test, Wilcoxon matched-pairs signed rank test and Kruskal-Wallis test, depending upon the suitability of the tests. Significance of normal data was examined with unpaired t-test, paired t-test, one-way ANOVA and two-way ANOVA (generalized linear mixed model) as required. Welch’s correction was applied to unpaired t-tests. Two-way ANOVA was always carried out with Geisser-Greenhouse correction. F values were rounded up to the closest integer. In the texts and figure legends, mean ± sem values are reported. Unless mentioned otherwise, the reported ‘n’ refers to the number of cells recorded from. Statistical significance notation were as follows: ns, p > 0.05; *, p= ≤ 0.05; **, p= ≤ 0.01; ***, p= ≤ 0.001; ****, p= ≤ 0.0001.

## Author approvals

All authors have seen and approved the manuscript. The manuscript has not been submitted or published elsewhere.

## Acknowledgements

We are grateful to Karin Dedek for helping us with retinal imaging, Denis Delkara and Olivier Marre for useful suggestions about marker gene targeting, and Mei Li for manufacturing AAV vectors. We would like to especially thank Gregory Schwartz for carefully reading our manuscript and for his helpful comments. This project was funded by NIH grants P30EY003176 and R01EY024334 to Richard H Kramer.

## Author contributions

Yadav SC, Nawy S and Kramer RH conceptualized the project; Yadav SC and Kramer RH designed the experiments; Yadav SC performed the electrophysiology experiments with help from Ganzen L and Nawy S; Man L helped with the acquisition of images; Ma H helped with the generation of cre-mice and gene editing; Yadav SC and Nawy S analyzed the data; Yadav SC and Kramer RH wrote the paper; Kramer RH supervised the project.

